# SurFlex Microscopy: Measuring Flexibility of Surface-Tethered Biomolecules

**DOI:** 10.1101/2024.12.13.628410

**Authors:** Aymeric Chorlay, Siddhansh Agarwal, Lena Blackmon, Daniel A. Fletcher

## Abstract

The flexibility of tethered molecules, such as those bound to biological membranes, is an important property that can influence molecular height, mobility, and accessibility. However, quantifying the flexibility of surface-tethered biomolecules in aqueous environments has been difficult due to a lack of experimental tools. Here we introduce SurFlex microscopy, a method based on fluorescence anisotropy that exploits the relationship between the conformational dynamics of a tethered molecule and the rotational diffusion of an attached fluorophore to extract information about molecular flexibility. By analyzing the polarization state of photons emitted after polarized excitation, we quantify apparent molecular flexibilities that include effects of tethering, self-interactions and buffer conditions. We first demonstrate the capabilities of SurFlex microscopy by measuring the flexibility of bilayer-tethered single-stranded DNA (ssDNA) of different lengths and nucleotide sequences. We find that sequence significantly impacts ssDNA flexibility, consistent with theoretical estimates, with weak intramolecular interactions in random sequences leading to higher apparent stiffness. Interestingly, we show that a pathological DNA sequence linked to Huntington’s disease exhibits unusual flexibility despite intramolecular interactions. We next extend SurFlex microscopy to live cells by measuring surface glycoprotein flexibility on red blood cells using fluorescent lectins. We show that trypsinization decreases glycan fluctuations, demonstrating that modifications to the cell surface can alter the flexibility of remaining surface molecules. SurFlex microscopy provides a new tool for quantifying molecular flexibility that can be used to study the role of tethered surface molecules in fundamental biological processes.

**Significance statement:** Biomolecules immobilized on one end play crucial roles in diverse cellular processes, from cell-cell signaling through surface receptors to the formation of DNA secondary structures. However, measuring biomolecular flexibility on surfaces has remained challenging. Here we present SurFlex microscopy, a technique that uses fluorescence anisotropy to quantify the flexibility of surface-anchored molecules. By analyzing the rotational dynamics of fluorophores attached to the ends of fluctuating biomolecules, SurFlex microscopy can be used to quantify persistence length. We demonstrate its capabilities by measuring sequence-dependent flexibility of DNA and crowding-dependent changes in glycan flexibility on native cell surfaces. This method opens new avenues for understanding how biomolecular flexibility influences key biological processes, such as those at cell surfaces during cell-cell contact formation and subsequent signaling.

## Introduction

The flexibility of polymer molecules in solution has been related to their dynamics through extensive theoretical and experimental work (1–6). However, many molecules exist in configurations where one end is tethered, introducing constraints that alter their dynamics, including reducing diffusion and increasing polymer-surface interactions (7–11). In biology, tethered molecules are very common, such as at the cell surface where tall mucins (e.g. MUC1) and short receptors (e.g. Fc receptors) extend above the plasma membrane by a few nanometers all the way up to hundreds of nanometers (12–15). The ability of membrane-tethered molecules to fluctuate is likely crucial to their biological functions, such as T cell receptors binding in trans to MHC molecules or cadherins oligomerizing in cis (16–18).

The factors affecting tethered polymer fluctuations are both intrinsic and extrinsic. The main intrinsic parameter is the molecule’s own chain flexibility, characterized by its bending energy, which determines the molecule’s ability to flex. Extrinsic factors from the molecular environment can also alter a polymer’s fluctuations through steric repulsions from crowded neighboring molecules, electrostatic interactions with the solvent and neighboring molecules, and specific binding interactions with other molecules, both tethered and soluble.

Several techniques have been used to investigate the fluctuations of molecules in solution, including fluorescence microscopy, atomic force microscopy (AFM), fluorescence resonance energy transfer (FRET), and optical trapping. The fluctuations of fluorescently labeled actin filaments were captured with microscopy and used to back out a bending rigidity of 7.3 10^−26^ N.m^2^ (19, 20). Atomic force microscopy (AFM) has been used to analyze the contours of free DNA on mica surfaces (21). FRET signal from fluorophores tethered to both ends of a DNA and peptidic molecule in solution has been used to quantify bending rigidity (22–25). For long tethered molecules like DNA and actin filaments, an optical trap was used to deduce molecular flexibility by measuring the force exerted by the molecule on a tethered microbead (26–28). Fluorescence polarization microscopy has been used to capture molecular orientation (29, 30), and fluorescence anisotropy can be used to measure the rotational motion of fluorescently labeled molecules and particles in solution (31, 32).

We wondered whether fluorescence anisotropy could be applied to molecules tethered to a surface rather than in solution. Instead of tracking a freely diffusing polymer with an attached fluorophore, we aimed to determine the bending rigidity of a surface-tethered polymer by measuring the rotational dynamics of its fluorescent label. While fluorophores could be added along the tethered polymer to assess local flexibility, we are particularly interested in the average chain flexibility based on the dynamics of a terminal fluorophore.

Here, we introduce SurFlex microscopy, a tool for measuring the flexibility of tethered biomolecules in aqueous environments. SurFlex microscopy is relatively simple to implement as it can be set up on a commercial confocal microscope with polarized excitation by imaging the orthogonal emission polarizations and computationally processing the polarization images. SurFlex microscopy leverages the interplay between a molecule’s conformational changes and the rotational motion of a fluorophore tag attached to its tip. By analyzing the polarization states of photons emitted by the tag, we can extract information about the molecule’s local rigidity from a theoretical model of a tethered polymer (see Supplementary Information). Unlike solution anisotropy measurements, anchoring the probed molecules on a surface constrains its orientational degrees of freedom and allows the characterization of short polymers at biologically relevant length scales (nanometers).

We demonstrate the capabilities of SurFlex microscopy by measuring the flexibility of short ssDNA strands anchored to a bilayer membrane. We find that the flexibility of ssDNA depends on both polymer length and nucleotide sequence. We show that weak intramolecular interactions, such as partial base pairings, considerably stiffen ssDNA, even when the interaction is only between a small number of nucleotides. Interestingly, we find that a pathological sequence linked to Huntington’s disease does not increase rigidity like other sequences, despite intra-chain base pairing. We then use SurFlex microscopy to characterize surface glycoproteins of red blood cells, showing that enzymatic removal of proteins reduces glycan fluctuations. Through quantitative measurements of DNA and glycocalyx flexibility, SurFlex microscopy opens new opportunities for studying fundamental biological processes that involve tethered molecules.

## Results

### Principle of SurFlex microscopy: determining the flexibility of surface-tethered molecules by analyzing the anisotropy of tip fluctuations

Biomolecules tethered to surfaces, such as the plasma membrane of a cell, fluctuate due to thermal motion with an amplitude limited by the rigidity of the molecule (Fig. 1A). To characterize the fluctuations of a molecule and thus determine its rigidity, SurFlex microscopy measures changes in the orientation of its tip. This measurement is achieved by attaching a fluorophore to the tip and exciting it with polarized light. Since light emitted from the fluorophore is also polarized, we can compare images of each orthogonal component to quantify the change in orientation over some lifetime.

**Fig. 1.**
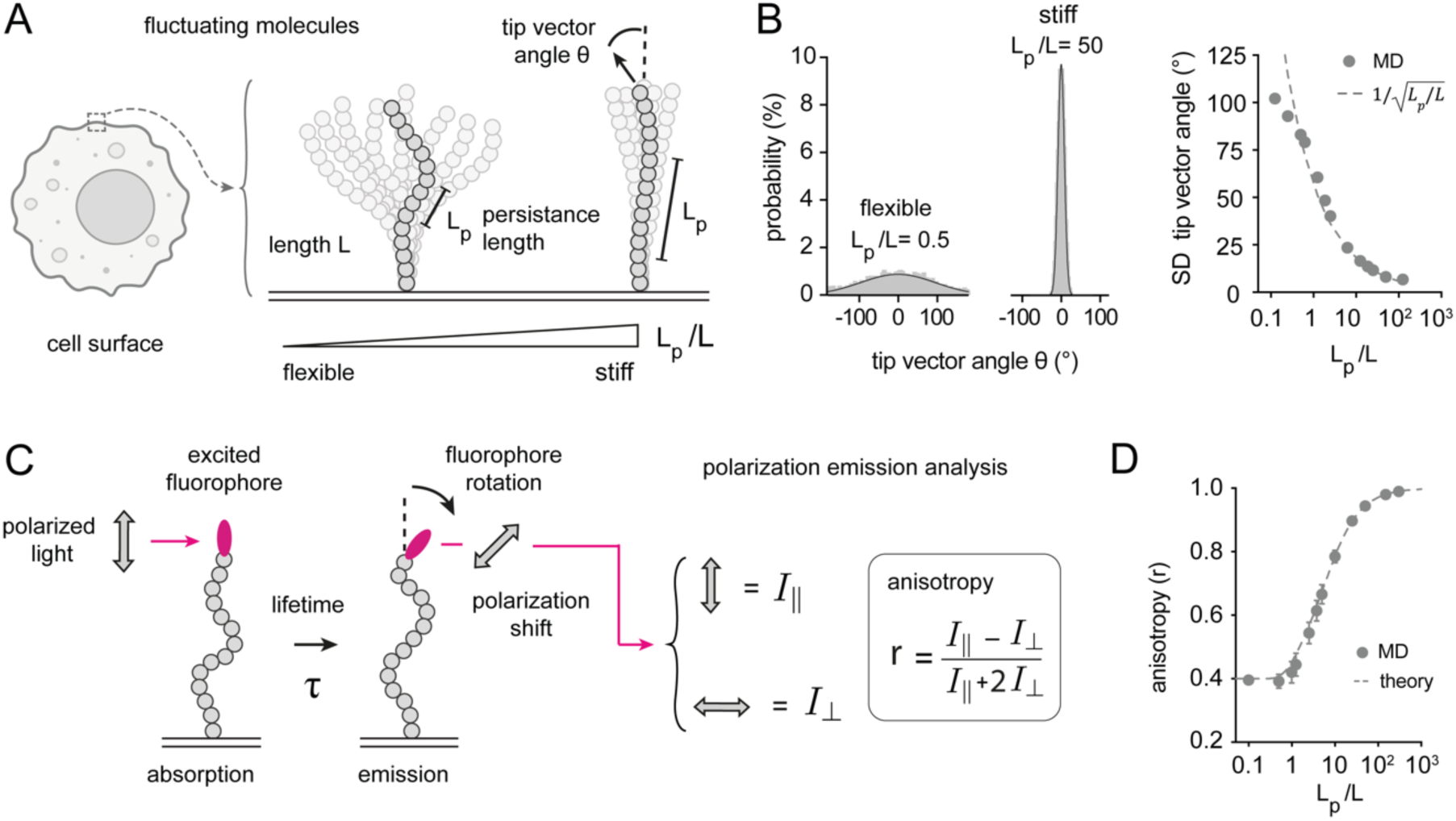
Tip fluctuations of tethered polymers are correlated with molecular flexibility. **(A)** Cell surface showing tethered molecules that extend above the membrane surface and fluctuate. Conformational dynamics of tethered molecules as a function of polymer stiffness, parameterized by the ratio of persistence length (*L_p_*) to contour length (*L*). The spread of angular distribution of the molecule’s end, reported by the tip tangent vector angle *θ* (TVA), decreases with increasing *L_p_*/*L*. **(B)** Quantitative analysis of molecule conformational states from molecular dynamics (MD) simulations. Left: Probability density distribution *p*(*θ*) of the TVA extracted from MD simulations for flexible (*L_p_*/*L* = 0.5) and stiff (*L_p_*/*L* = 50) polymer; solid curves represent theoretical prediction from equation (1). Right: Standard deviation *σ* of *p*(*θ*) as a function of *L_p_*/*L*. MD data points lie on top of *σ* = 1/.*L_p_*/*L* (dashed curve) scaling from the simple WLC model for *L_p_*/*L* >> 1. **(C)** Fluorescence anisotropy measurement for polymer-tethered dipoles: Photoselection involves excitation of dipoles aligned with the electric field vector of incident linearly polarized light. During the excited state lifetime (τ), the dipole attached to the polymer undergoes rotational diffusion driven by the polymer’s conformational fluctuations. Consequently, the emitted light’s polarization axis deviates from the excitation orientation. Polarization-resolved detection of emitted photons along axes parallel (I_ǁ_) and perpendicular (I_⊥_) to the excitation polarization enables quantification of the dipole’s rotational degrees of freedom. The relative intensities of I_ǁ_ and I_⊥_ are functions of both orientational distribution and rotational diffusion: a narrow distribution with minimal rotation results in I_ǁ_>> I_⊥_, while a broad distribution and/or significant reorientation leads to increased I_⊥_. Anisotropy (r) definition based on polarized emission components I_ǁ_ and I_⊥_. **(D)** MD-simulated anisotropy (r) values for a 14 unit long polymer molecule as a function of *L_p_*/*L*. Results demonstrate anisotropy’s sensitivity to stiffness (*L_p_*/*L*), data closely matching theoretical predictions.

To validate this approach, we first confirmed that fluctuations of the molecule’s tip correlate with chain stiffness. The persistence length (*L_p_*) of a molecule is often used to quantify its rigidity and is related to the bending energy (*κ*) by the equation *L_p_* = *κ*/*k*_*B*_*T*, where *k_B_* is the Boltzmann constant and T is temperature (33). Smaller *L_p_* values correspond to more flexible, coil-like states, and larger values indicate stiffer, more rod-like conformations (Fig. 1A). Tip fluctuations are reported by the tip vector angle *θ* (TVA), defined as the angle between the molecule’s tip vector and the surface normal (Fig. 1A, top right). We carried out molecular dynamics simulations (MD) modeling the tethered polymer as a Worm-Like Chain (WLC). The key dimensionless parameter used to characterize the molecular fluctuations is the ratio *L_p_*/*L*, where *L* is the total length of the chain. This ratio encapsulates the competition between bending energy and entropy, with molecules of greater contour length exhibiting larger fluctuations for a given persistence length. The *L_p_*/*L* ratio determines chain behavior: >1 (stiff), ∼1 (semi-flexible), <1 (flexible). Simulations show that the TVA distribution narrows markedly as *L_p_*/*L* increases (Fig. 1B, left). The probability distribution of the TVA is well-approximated by the expression (34, 35):

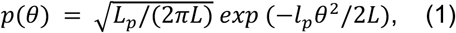

plotted as solid curves in Fig. 1B, left. The spread of the distribution of TVA is characterized by the standard deviation *σ* = 1/.*L_p_*/*L*, agreeing with MD simulations for *L_p_*/*L* > 1 (Fig. 1B right, see SI Appendix for details). These results validate the TVA as an effective indicator of molecular fluctuations.

To characterize the orientation distribution of the polymer’s TVA, we attach a fluorophore to the polymer’s free end, which acts as a sensitive probe for the polymer’s tip orientation (Fig. 1C). When illuminated with linearly polarized light, the fluorophore preferentially becomes excited if it aligns with the light’s polarization vector (36), with an excitation cos^2^ probability proportional to *P*(*θ*_*a*_) ∝ *cos*^2^(*θ*_*a*_), where *θ*_*a*_ is the angle between the fluorophore’s absorption dipole and the electric field vector of the polarized light. Following excitation, the fluorophore rotates along with the polymer tip without emitting light. At the end of its fluorescent lifetime *τ* (∼ns), the fluorophore emits light polarized along its new orientation. During the millisecond timescale of the measurement, the fluorophore undergoes many cycles of absorption and emission, allowing it to probe the angular rotation of the polymer tip approximately a million times.

To quantify the orientation and rotational freedom of the fluorophore-labeled polymer tip, we measure fluorescence anisotropy (*r*), which is the normalized difference between the parallel *I*_∥_ and perpendicular *I*_D_ polarization components of emitted light relative to the excitation polarization:

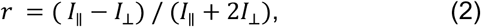

In the stiff limit (*L_p_*/*L* > 1), the polymer’s bending rigidity restricts the angular space available to the tip (Fig. 1B). This spatial restriction limits rotational diffusion around the polymer axis, causing the emitted light to maintain strong parallel polarization (*I*_∥_). Consequently, the anisotropy values remain high, reporting the narrow orientation distribution of the tethered molecule. Conversely, flexible molecules (*L_p_*/*L* < 1) allow the fluorophore to explore a broader angular range through rotational diffusion (Fig. 1B). This increased orientational freedom enhances *I*_D_ relative to *I*_∥_, resulting in lower *r*. As polymer flexibility increases further, the anisotropy approaches a limit as the fluorophore achieves nearly unrestricted rotational diffusion.

Our theoretical model predicts that anisotropy increases with polymer stiffness, a relationship validated by both theoretical calculations and molecular dynamics simulations (detailed in SI appendix). As demonstrated in Fig. 1D, this correlation is robust in *L_p_*/*L* > 1 regime, where *r* serves as a direct measure of polymer stiffness, indicating that an anisotropy measurement with a fluorophore at the molecule’s tip allows us to quantify flexibility of the tethered molecule.

### Measurement of molecular flexibility with SurFlex microscopy

To test flexibility measurements of tethered molecules experimentally, we attached fluorescently labeled molecules to the surface of giant unilamellar vesicles (GUVs) (Fig. 2A). These vesicles are imaged using a spinning disk confocal microscope, modified to excite samples with linearly polarized light. The emitted light is collected by an objective lens and directed towards a polarizing beam splitter, which separates the vertically and horizontally polarized components. These components are then filtered by one of two linear polarizers that are oriented orthogonally (Fig. 2A). A camera simultaneously captures the two corresponding images, measuring the intensities I_ǁ_ and I_⊥_, which are then used to calculate anisotropy and determine molecular rigidity (*L_p_*/*L*).

**Fig. 2.**
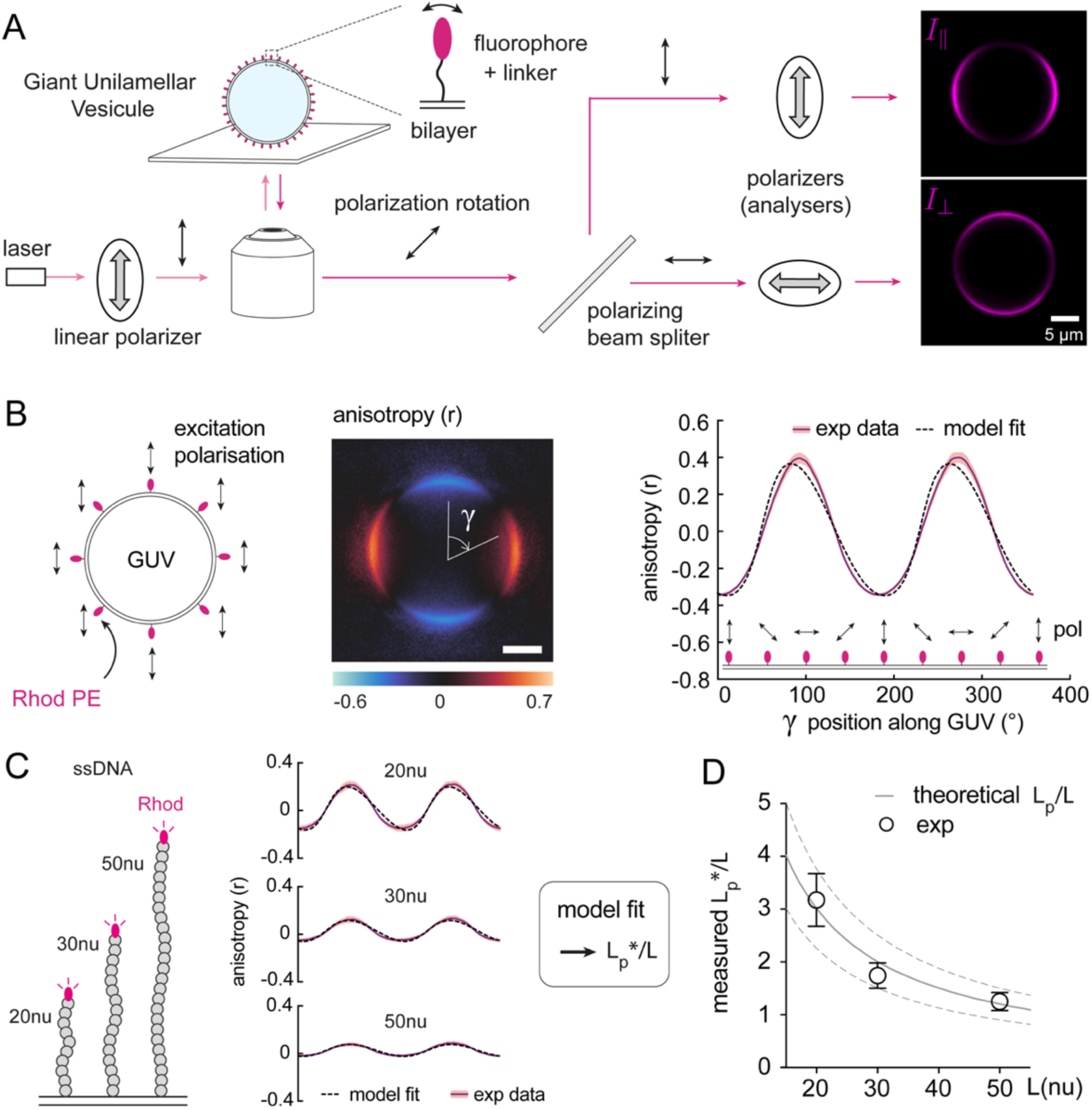
Fluorescence anisotropy measurements of tethered ssDNA tip fluctuations enable quantification of flexibility. **(A).** Left: Spinning disk confocal microscope system to measure anisotropy of tethered molecules to the bilayer of a giant unilamellar vesicle (GUV). Laser light source is filtered by a polarizer to obtain linearly polarized light that excites the fluorophore attached to the end of the molecule of interest anchored to the GUV surface. The fluorophore absorbs the light when its dipole aligns with the polarization. The fluorophore changes orientation due to thermal fluctuations during its excited state lifetime. It then emits light polarized accordant with its new orientation. This light is collected by the objective and decomposed into two orthogonal linear polarizations by the polarizing beam splitter. Each component is further filtered by a polarizer and imaged simultaneously by the camera. Right: image obtained from a GUV displaying fluorophores attached by a short linker to its bilayer (lissamine Rhodamine–Phosphatidylethanolamine termed Rhod PE). Top image: light intensity polarized parallel to the excitation polarization (I_ǁ_); Bottom image: light intensity polarized perpendicular to the excitation direction (I_⊥_). **(B) Left:** Anisotropy image calculated from registered images of I_ǁ_ and I_⊥_ for Rhod PE molecules distributed radially on the surface of a GUV. Note that the value of the anisotropy varies along the GUV contour, as the molecules do not all have the same orientation relative to the polarization of the excitation light. **Right:** Line scan of the anisotropy as a function of the angle *γ* reporting the position along the bilayer. Experimental data are mean values (purple solid line) and standard deviation (light pink shaded area) of n > 30 line scans performed on independent GUVs. The sketch below the curve shows the relative orientation of the Rhod PE molecules and the excitation polarization. The anisotropy is greatest when the polarization and the Rhod PE molecule are perpendicular, suggesting that the Rhodamine dipole is oriented tangentially to the bilayer. The data obtained are fitted by a model (r^2^=0.97) which allows us to determine the intrinsic parameters of the fluorophore (for full model description and fitting procedure, see *SI appendix,* supporting text). **(C)** Anisotropy line scan of GUV-tethered single strand DNA (ssDNA) of variable length (L). Experimental data are mean values and standard deviations (respectively purple solid line and light pink shaded area) of n > 30 line scans performed on independent GUVs. Experimental data are fitted using the Rhodamine parameters obtained in B and by adjusting the *L_p_*/*L* value of the model (20nu r^2^=0.97; 30nu r^2^=0.96; 50nu r^2^=0.97; for full model description and fitting procedure, see methods and *SI appendix* supporting text). **(D)** Measured apparent *L_p_*\*/*L* values, obtained by fitting each GUV line scan from C, as a function of ssDNA length in base pair length units. The error bar indicates SD. Solid gray line indicates the theoretical *L_p_*/*L* fit (r^2^=0.95), dashed line indicates the 95 percent confidence interval.

We used Lissamine Rhodamine conjugated to a PE lipid (Rhod PE) present at 1% in the GUV membrane to obtain I_ǁ_ and I_⊥_ images of a short tethered molecule (Fig. 2A, right). Each pixel (0.2 µm x 0.2 µm) captures the average polarization of approximately 600 fluorophores. Moreover, during a 200 ms exposure, each fluorophore, with a lifetime of 2.6 ns (measured by FLIM, Leica Stellaris), is probed around a million times. As a result, the intensity measured for each polarization component is a population average that is measured many times, providing a statistically robust measure of local polarization. The I_ǁ_ image reveals an asymmetric intensity distribution, peaking at the lateral poles of the vesicle. This suggests a preferentially tangential orientation of the Rhod PE dipole with respect to the membrane, which is important to consider while computing anisotropy (see *SI appendix* for details).

Images I_ǁ_ and I_⊥_ are then used to compute the anisotropy (r) of Rhod PE molecules. A custom MATLAB program detects the GUVs, registers the I_ǁ_ and I_⊥_ images, then calculates the anisotropy pixel by pixel (see Materials and Methods). The program generates a map of the anisotropy of Rhod PE on the vesicle surface (Fig. 2B, center). This anisotropy map is then analyzed by a line scan along the contour of the GUV, revealing anisotropy values as a function of angle *γ*, which defines the position along the GUV perimeter (Fig. 2B, right).

Notably, we observe that anisotropy varies cyclically as a function of position, with a minimum at *γ* = 0° and maximum at *γ* = 90°. One might expect constant anisotropy along the vesicle contour, since there is no reason to expect molecules on one part of the GUV to have a different flexibility from another. While this is correct, the variation is due to the geometry of the system. Although the polarization of the excitation light is spatially uniform, the Rhod PE molecules are oriented radially with respect to the vesicle surface (Fig. 2B, left). Consequently, the angle between the axis of each molecule and the direction of excitation light polarization changes as a function of position on the vesicle (see cartoon above *γ* axis, Fig. 2B, right). Thus, the variation in anisotropy reflects the variable alignment of the molecules with the incident polarization and not any heterogeneity in their stiffness.

Typically, to extract the molecular stiffness (*L_p_*/*L*) from measured anisotropy, we must also include crucial fluorophore-specific parameters. The angle between the fluorophore’s absorption and emission dipoles (β) – a property arising from the fluorophore’s electronic structure – and the fluorophore’s orientation relative to the polymer backbone (α) critically influence the anisotropy measurements. Our model combines the fluorophore’s orientation angles (α, β) with the polymer’s conformational ensemble to establish a direct relationship between measured anisotropy and persistence length (see *SI appendix,* supporting text for derivation).

Rhod PE is very short and therefore has a higher stiffness value (*L_p_*/*L* >> 1) resulting in an anisotropy signal of high amplitude. This enabled us to measure fluorophore-specific parameters needed for quantifying the flexibility of less stiff molecules labeled at the tip with Rhodamine. The absorption and emission dipoles of Rhodamine have already been characterized (37, 38) and found to differ by 11.9° (β = 11.9°; see *SI appendix,* supporting text for details). Using this value, we determined that the fluorophore attaches nearly perpendicular to the polymer axis (α = 82°). These fluorophore-specific parameters are included in the theoretical model, enabling us to estimate the rigidity of the imaged molecules. Since both internal and environmental constraints affect the molecule’s fluctuations, we use the apparent persistence length (*L_p_*\*) as the measured value to distinguish from persistence length in the absence of other effects (*L_p_*).

Next, to validate that the SurFlex microscopy is capable of quantifying molecular flexibility beyond the rigid model regime, we quantified the anisotropy of ssDNA oligomers of varying lengths (20, 30 and 50 nucleotides), which should show different *L_p_*\*/*L* due to their different lengths. These oligomers are labeled at one end with a Rhodamine fluorophore and anchored to the GUV surface by a cholesterol moiety (Fig. 2C). We find that the amplitude of the anisotropy profiles decreases with increasing oligomer length, indicating increasing fluctuation and decreasing stiffness. By only tuning *L_p_*\*/*L*, our model fits the anisotropy profiles very well, enabling us to extract the apparent stiffness of each oligomer. As expected, the measured stiffness decreases with oligomer length, following a 1/L trend (measured stiffness *L_p_*\*/*L*, Fig. 2D).

### Measurement of ssDNA persistence length and the effect of nucleotide sequence

We next used SurFlex microscopy to evaluate the influence of nucleotide sequences on the persistence length of ssDNA. We designed two ssDNA oligomers of fixed 21 nucleotide length: one composed solely of adenosines (polyA), the other of cytosines (polyC). After anchoring them to the surface of GUVs, we measured anisotropy and applied our model to extract *L_p_*\*/*L*. Using the number of nucleotides and their average size of 0.5 nm (39–42), we calculated the total length (*L*) of the strands (10 nm) and computed the measured apparent persistence length for each oligomer. We found that polyA has a higher apparent persistence length than polyC (Fig. 3A). This is in agreement with theoretical predictions (43) and experimental characterization (23, 39), suggesting that adenosines, which have a larger double-ring nitrogenous base structure compared to single-ring cytosines, limit the bending motions between segments of the ssDNA chain, thereby increasing its rigidity.

**Fig. 3.**
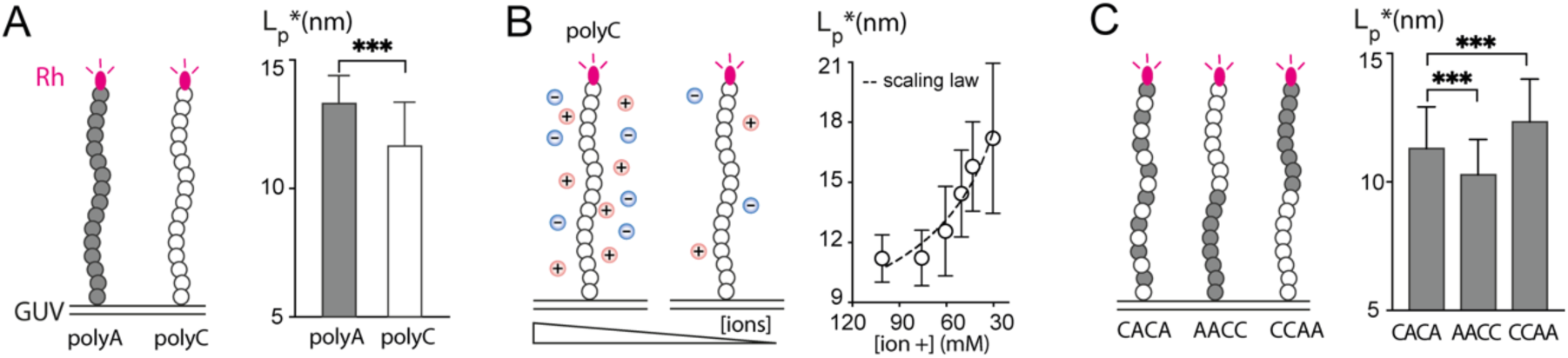
Flexibility of ssDNA is sequence dependent. **(A).** Measurement of the apparent persistence length (*L_p_*\*) of ssDNA oligomers of identical length but different sequences – polyA (21 adenine) and polyC (21 cytosine)– showing a significant difference (student test p< 0.0001). Data obtained by fitting the anisotropy line scan of n > 60 independent GUVs for each condition. Error bar indicates standard deviation. **(B).** Measurement of *L_p_*\* for polyC (21 cytosine) ssDNA oligomers while decreasing the ion concentration in solution by adding MQ water. Data obtained by fitting the anisotropy line scan of n > 20 independent GUVs for each condition. Error bar indicates standard deviation. Dashed line is a scaling law: *L_p_* ∗= *L_p_* + *A*√*C where L_p_* ∗ is the apparent persistence length, A and *L_p_* are fitting parameters and C is the concentration of positive ions. r^2^=0.94. A=2.6nm. M^1/2^ and *L_p_* =2.4nm. **(C).** Measured *L_p_*\* for 21 nucleotide long ssDNA oligomers with different combinations of Adenine (A) and Cytosine (C): CACA: alternating C and A; AACC: 11 A followed by 10 A; CCAA: 11 C followed by 10 A. CACA shows a statistically significant difference compared to AACC and CCAA (respectively Mann-Whitney test p< 0.0001 and student test p< 0.0001). Data obtained by fitting the anisotropy line scan of n > 60 independent GUVs for each condition. Error bar indicates standard deviation.

Based on the SurFlex microscopy measurements, we calculated an *L_p_*\* of approximately 13 nm, an apparent persistence length of the ssDNA oligomer, encompassing all constraints experienced by the chain in addition to its intrinsic rigidity. The intrinsic persistence length values reported in the literature, which vary greatly depending on the techniques used, range from 0.7 to 7 nm (28, 39, 44–49). This suggests that the ssDNA strands we measured are subject to additional constraints that increase their apparent rigidity. We hypothesized that this phenomenon could be mainly due to repulsive electrostatic interactions between negatively charged phosphate groups. These repulsions induce long-range correlations in the polymer conformation, reducing conformational fluctuations and thus increasing the apparent persistence length (50–53). As the intensity of these repulsions directly depends on the concentration of positive ions, we postulated that a decrease in the ionic concentration of the medium would result in an increase in the apparent persistence length (*L_p_*\*).

To test this hypothesis, we measured the persistence length of a cytosine oligomer (polyC) while gradually diluting the solution with water, thereby progressively reducing the ionic concentration (Fig. 3B). Decreasing the positive ion concentration indeed causes an increase in the measured persistence length, in agreement with previous work (50, 51). This increase in persistence length as a function of concentration follows a power-law scaling with an exponent of 1/2, consistent with the scaling law postulated by Joanny et al (54) (see *SI appendix,* Fig. S4). By fitting these results with the scaling law derived from Joanny et al. *L_p_* ∗= *L_p_* + *A*√*C*, we estimate the intrinsic persistence length (*L_p_*) of the polyC chain to be around 2.4 nm which falls within the range of values reported in the literature (28, 39, 44–49).

Having observed that ssDNA strands of different nucleotide compositions exhibit distinct persistence lengths, we sought to determine whether mixing these two types of nucleotides by alternating them (CACA) would yield an intermediate *L_p_*\*. The measurement of the apparent persistence length of this mixed oligomer reveals a value close to that of polyC, significantly lower than that of polyA (Fig. 3C). This suggests that the introduction of cytosines between adenosines, due to their smaller size, increases the flexibility of the chain. These results suggest that the flexibility of the A-C-A linkage is comparable to that of the C-C-C linkage.

Next, we examined the effect of spatial segregation of nucleotides while maintaining the same overall composition of adenosines (A) and cytosines (C). For this purpose, we designed two new oligomers: (i) a polyA sequence close to the anchoring point, followed by a polyC sequence carrying the fluorophore AACC and (ii) a polyC sequence close to the anchoring point, followed by a polyA sequence carrying the fluorophore CCAA (Fig. 3C). Surprisingly, we measured significantly different *L_p_*\* for these two sequences. The CCAA oligomer proved more rigid than the alternating sequence CACA, while the AACC oligomer was more flexible. These results suggest that the nucleotide sequence directly influences the measured flexibility, beyond the simple composition, opening new avenues for understanding the relationship between nucleotide sequence and mechanical properties of nucleic acids.

### SurFlex microscopy shows that the apparent persistence length of ssDNA strongly depends on weak self interactions

Although ssDNA sequences composed solely of A’s and C’s are a convenient model to study ssDNAs that do not interact with themselves, they do not represent the reality of most DNA sequences present in nature, which make use of all four bases. We therefore wanted to know how the persistence length of ssDNA is affected by a mix of nucleotides. To this end, we randomly generated a collection of 21 nucleotides composed of A, T, G, and C with a constant 30% GC content. From this collection, we synthesized three different ssDNA oligomers (named seq#1, seq#2, and seq#3) and measured their apparent persistence lengths (Fig. 4A). Interestingly, all the oligomers had an apparent persistence length greater than that of polyC, with seq#3 having an apparent persistence length four times higher (46 nm). How can we explain these unexpected values of apparent rigidity? Before synthesizing the random oligomers, we verified that they did not form hairpin structures, though the possibility remains that the nucleotides could transiently interact with each other by forming base pairs that could rigidify the oligomers.

**Fig. 4.**
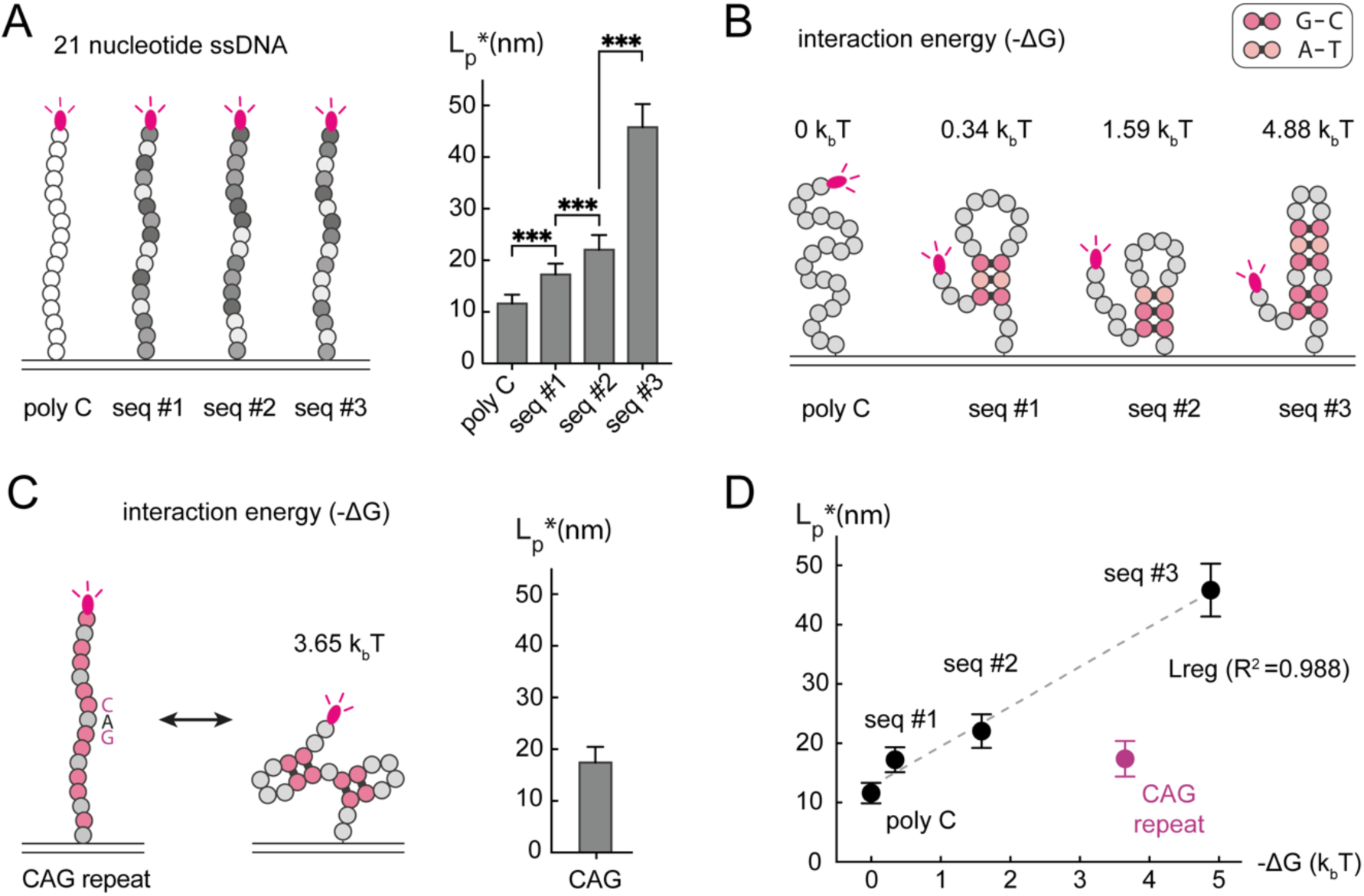
Apparent persistence length of ssDNA depends strongly on weak self-interactions. (**A).** Measurement of apparent persistence length (*L_p_*\*) of ssDNA with a random permutation of Adenine, Cytosine, Guanine, and Thymine, keeping the nucleotide content constant (GC content 30%). The *L_p_*\* of a 21-cytosine oligomer is presented as a reference. Data obtained by fitting anisotropy line-scans of n > 30 independent GUVs for each condition. P values are < 0.0001 based on Mann-Whitney test for PolyC-seq#1 and seq#1-seq#2 comparisons. P values are < 0.0001 based on Welch’s t test for seq#2 to seq#3 comparison. Error bar indicates standard deviation. (**B).** Most probable structure obtained with NuPack 4.0 software for the ssDNA measured in A. The associated interaction energy (-ΔG) is calculated by weighting the energies of the three most prevalent structures by their respective probabilities. For the complete set of structures and probabilities, see *SI appendix* Fig. S5. **(C).** Left: Measurement of *L_p_*\* for a ssDNA composed of seven repeats of the Cytosine Adenine Guanine (CAG) sequence, which is a marker associated with Huntington’s disease. Data obtained by fitting anisotropy line-scans of n = 30 independent GUVs for each condition. Error bar indicates standard deviation. Right: Most probable structures and associated interaction energy (-ΔG) obtained with NuPack 4.0 software. For all structures and associated probabilities see *SI appendix* Fig. S5. **(D).** *L_p_*\* is plotted against the folding energies associated with each sequence described in A and C. For random ssDNA fragments and poly C, measured *L_p_*\* increases with folding energy (-ΔG). CAG repeat does not seem to follow this pattern. Dashed line is a linear regression (R^2^= 0.988) of the data points, excluding CAG repeats. Error bar indicates standard deviation.

We investigated the potential for sequence interactions using the NuPack 4.0 software and obtained a set of transient structures for each oligomer, as well as the associated energies (*SI appendix* Fig. S5). From these data, we calculated for each oligomer an interaction energy (-ΔG) that corresponds to the sum of the energies of each structure, weighted by their respective probability. The higher the folding energy, the longer-lived the oligomer structures are likely to be. Fig. 4B represents the most probable structures and their associated interaction energies (-ΔG). We noticed that the most stable structure, seq#3, which exhibits the highest apparent persistence length (*L_p_*\*), has a folding energy (-ΔG) of 4.88 k_b_T. For comparison, a hairpin structure has an energy of −ΔG = 17 k_b_T (*SI appendix* Fig. S5).

We next wanted to investigate whether there was a connection between interaction energy and persistence length for a real gene sequence. We chose to study the repeated CAG motif that is a marker for Huntington’s disease. A proliferation of CAG repeats in the gene coding for huntingtin protein causes the formation of a defective protein that accumulates in neurons (55, 56). We synthesized an oligomer presenting seven CAG repeats and measured its apparent persistence length, which turned out to be around 17 nm, close to that of seq#1 (Fig. 4C, left). However, calculating its expected sequence interactions, we found significant structuring, close to that of the seq#3 oligomer (Fig. 4C, right).

To compare these observations, we plotted the measured persistence length values as a function of the folding energy (-ΔG) (Fig. 4D). For the random sequences, we observed a positive correlation between persistence length and folding energies, except for the CAG repeat, which maintains high flexibility despite its structuring. It is possible that repetition of the CAG sequence allows the formation of relatively stable base pairs, distributed homogeneously, thus not overly restricting the rotational degrees of freedom of the molecule. It is interesting to note that this particular disease-associated sequence does not follow the behavior of the other sequences, highlighting the complexity of structure-flexibility relationships for nucleic acids. These observations demonstrate the sensitivity of the SurFlex microscopy in detecting subtle intramolecular interactions within ssDNA chains, which contribute to increasing their apparent rigidity.

### SurFlex microscopy measures dynamic changes in fluctuations of glycosylated molecules on red blood cells during trypsinization

To demonstrate SurFlex microscopy on live cells, we studied the fluctuation of red blood cell surface glycosylated molecules. We labeled these molecules with fluorescent lectins and measured anisotropy, with the goal of quantifying their fluctuations and assessing how they would be affected by trypsin-induced removal of surface proteins.

We began by labeling the RBCs’ glycosylated surface molecules using RCA-I, a lectin tagged with fluorescein, which binds to terminal galactose (57, 58). To measure anisotropy, we created spherical RBCs by swelling the cells in a hypotonic solution (Fig. 5A). The measured anisotropy values (Fig. 5B, 0 min) indicate significant molecular flexibility, though direct comparison with ssDNA measurements is precluded by differences in fluorophore properties. Trypsinization, which cleaves surface proteins at lysines and arginines, causes an increase in the amplitude of the anisotropy measured on RBCs (Fig. 5B), suggesting a reduction in the fluctuations of glycosylated molecules.

**Fig. 5.**
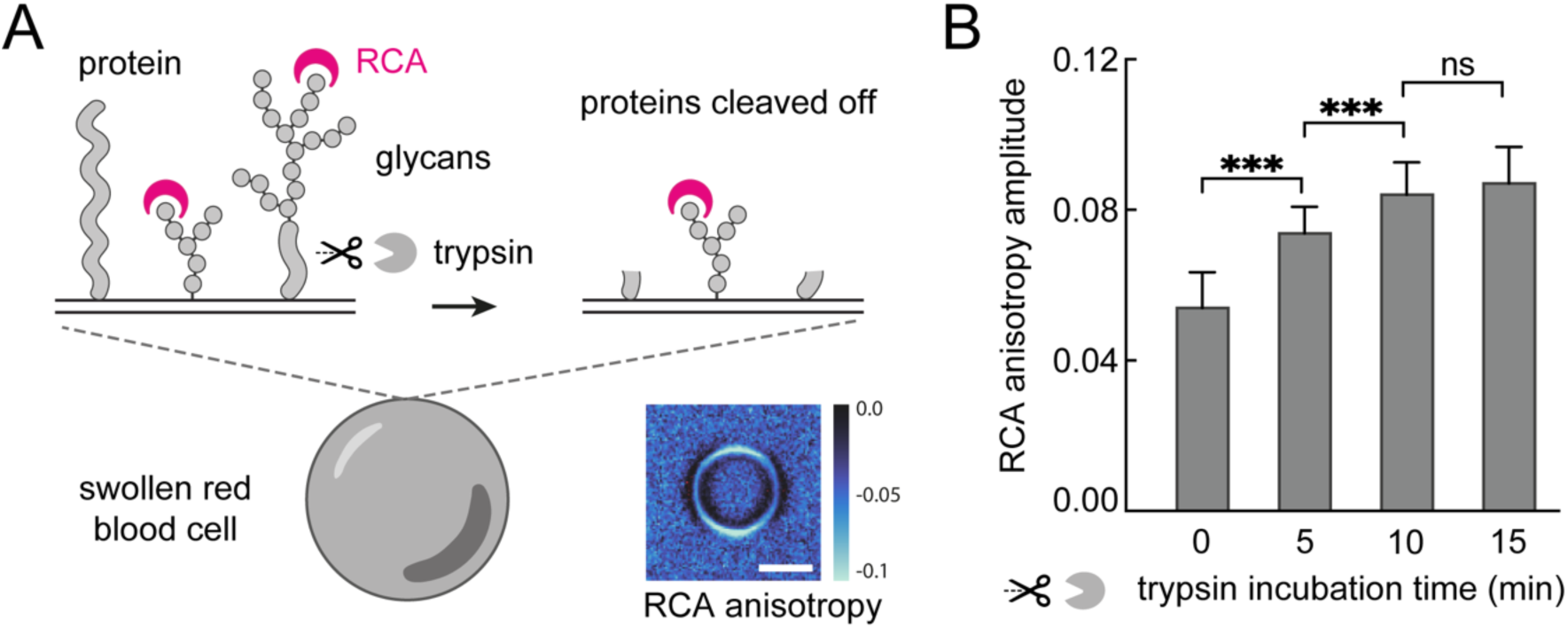
Fluctuation measurements of glycosylated molecules on red blood cells. **(A)** Red blood cells (RBCs) are osmotically swollen to become spherical. The terminal galactose residues of glycosylated molecules on the RBC surface are labeled with fluorescein-conjugated RCA lectin. Surface proteins on the RBCs are cleaved by incubation with trypsin for varying times. Anisotropy image of RCA lectin on the surface of a RBC. Scale bar is 5μm. **(B)** Anisotropy amplitude of fluorescein-labeled RCA lectin on the surface of RBCs incubated with trypsin for 0 to 15 minutes. Data were obtained by measuring the anisotropy amplitude of n=30 independent RBCs for each condition. P-values are < 0.0001 based on t-tests for the comparisons between 0 and 5 minutes, and 5 and 10 minutes. There is no statistical difference between the 10 min to 15 min conditions (P-value 0.196). Error bar indicates standard deviation.

Our interpretation is that the rather bulky RCA lectins (120 kDa) (58) first access the tall, highly fluctuating glycosylated molecules with low measured anisotropy. When trypsin cleaves off proteins, reducing their length, the remaining galactose residues on shorter surface molecules fluctuate less, resulting in higher anisotropy. It is also possible that protein cleavage reduces surface density, allowing the RCA lectins to access smaller sugars (on galactosylated lipids) that fluctuate less due to their size. This interpretation is supported by measurements of RCA binding level to the RBCs’ surface, which show that RCA binding first increases after 5 minutes of trypsin treatment, then decreases with prolonged exposure (*SI appendix* Fig. S6). This pattern likely results from two processes. First, trypsin removes dense surface proteins that initially block RCA access to surface galactose. Subsequently, continued trypsinization removes glycosylated surface proteins, reducing available RCA binding sites but allowing access to smaller and less flexible sugars. These data demonstrate that SurFlex microscopy is capable of directly measuring changes in biomolecule fluctuations on cells.

## Discussion

The flexibility of biomolecules is important for their biological activity in many contexts. For example, DNA must interact with numerous partner proteins, such as enzymes involved in replication, repair, or gene expression. These interactions often require DNA to bend and fold around the proteins to form complexes. Thus, being able to characterize the flexibility of DNA at the scale of a few nucleotides, particularly in regulatory regions and protein binding sites, appears to be a major challenge to understanding the molecular mechanisms governing the access of genetic information. Moreover, a precise measurement of DNA flexibility could be useful for designing DNA nanotechnology, such as origami for drug delivery and hairpin probes for molecular detection.

We introduce SurFlex microscopy as a new approach to directly measure the flexibility of single-stranded biomolecules. By combining fluorescence anisotropy and theoretical modeling, we can quantify flexibility of surface-tethered molecules. Our study reveals significant differences in apparent persistence lengths between homopolymers of ssDNA, reflecting the impact of nucleotide composition on molecular rigidity, as predicted by previous studies (23, 39, 43). We measure a persistence length difference (*ΔL_p_*) of approximately 2 nm between polyA and polyC. If we consider the interaction of these ssDNA strands with a typical 50 kDa protein having a typical radius of 2 nm (59), the bending energy associated with folding the DNA strand around the protein scales as *U* ∼ *L_p_L*/2*R*^2^ in k_b_T units. With a persistence length difference *ΔL_p_* ∼ 2*nm*, this would mean a supplemental energy cost of *ΔU* ∼ 3 *k*_*b*_*T* to fold polyA compared to polyC, which could have a significant impact on the energetics of protein-DNA interactions, which are on the scale of ∼ 10 *k*_*b*_*T* (60, 61). If the nucleotide type affects the flexibility of single-stranded DNA, it is possible that chemical modifications could also influence its flexibility. Changes in DNA flexibility due to modifications could potentially affect processes like transcription, replication, and repair. For example, DNA methylation that influences gene expression by modifying transcription factor binding could possibly be explained by a change in DNA flexibility that would affect the accessibility of binding sites or the stability of the protein-DNA complex.

Our findings show that both the composition and sequence order of nucleotides significantly modulate ssDNA stiffness, with the terminal sequence playing a dominant role in determining apparent flexibility. SurFlex microscopy primarily reports the flexibility of the region where the fluorophore is attached, as evidenced by the striking correlation between the apparent persistence length (*L_p_*\*) of heteropolymers and their terminal segments. For instance, sequences with adenine at the free end (e.g., CCAA) exhibited a higher *L_p_*\* than those with cytosine at the free end (e.g., AACC), mirroring the trend observed in poly(A) and poly(C) homopolymers (Figs. 3A,C). The higher *L_p_*\* of A-terminated constructs likely results from adenine’s stronger base stacking interactions (62), higher backbone charge density, and smaller hydration shell (63). While the terminal segment dominates the measured flexibility, the proximal segment exerts a secondary influence on the overall conformation, subtly affecting the distal region’s behavior. This is exemplified by the intermediate *L_p_*\* of alternating sequences like CACA. These sequence-dependent variations in stiffness reveal the intricate interplay between nucleotide arrangement and ssDNA mechanical properties, offering new insights into nucleic acid behavior and highlighting the potential for fine-tuning ssDNA flexibility through strategic sequence design.

More unexpectedly, we demonstrate that random sequences can exhibit increased apparent rigidity due to weak self-interactions, such as partial base-pairing. The importance of these intramolecular interactions appears in the correlation between apparent persistence length and folding energy of the oligomers (Fig. 4D). However, this trend is not followed by ssDNA with CAG repeats, which we found to be unusually flexible despite their structuring, showing the physical complexity of nucleotide interactions. In some instances, it is possible that a structured oligonucleotide appears to be flexible but is, in actuality, not. For example, a molecule may have a structure that allows oscillations between two distinct states that are both rigid. Our measurements will likely classify this molecule as flexible, highlighting the limitations of describing rigidity purely in terms of the worm-like chain model’s internal stresses. It would be more accurate to describe it in terms of fluctuations between different states. For complex biomolecules, especially those on cell surfaces, both structural states and dynamic transitions must be considered.

SurFlex microscopy can also be used to directly measure flexibility of molecules tethered to the cell surface. We used red blood cells and demonstrated that trypsinization reduced the fluctuations of glycosylated molecules on their surface, likely by exposing shorter, less fluctuating molecules to our measurement. However, our labeling agent (lectin RCA) indiscriminately labels all accessible molecules with terminal galactose moieties (57, 58). Our measurement is therefore an average of the fluctuations of all molecules with galactose groups. To refine the results obtained with our technique, it is necessary to use fluorescent labeling methods that selectively target molecules of interest. Furthermore, to accurately report fluctuations in the target molecule, it is essential to label them with fluorophores that are smaller than the target. If the marker is too large, its own fluctuations will contribute to the measurement, while its inertia will dampen fluctuations of the target molecule. For work involving proteins, labeling techniques based on small molecules are preferable, such as FlAsH, which involves a small molecule fluorophore that binds to a specific short amino acid sequence engineered into the target protein (65). Through careful labeling strategies and appropriate model to interpret fluctuations, SurFlex microscopy can capture accurate measurements of biomolecule flexibility.

In conclusion, this study introduces SurFlex microscopy as a new method for quantifying tethered polymer flexibility. Our experimental measurements of ssDNA demonstrate the impact of DNA sequence on flexibility and reveal unexpected behaviors that illustrate the complex interplay between primary structure and mechanical properties. Beyond DNA, SurFlex microscopy can be used to study the mechanical properties of other complex biomolecules, both on synthetic membranes and on the surface of living cells.

## Materials & Methods

### Microscope used for anisotropy measurement

SurFlex was implemented on a Nikon Ti-2 microscope equipped with a spinning disk (Yokogawa CSU-X1). A linear polarizer (LpVis-100A from Thorlabs) was placed in the excitation laser path. Polarized images were captured by inserting a dual image module (Hamamatsu) equipped with a polarizing beam splitter (Moxtek FBF04C) and two orthogonal polarizers (LpVis-100A from Thorlabs) between the emission filter and the camera.

### Preparation of ssDNA samples for measurement

#### Preparation of Giant Unilamellar Vesicles (GUVs)

The lipid mix used to form the lipid membrane of the GUVs was kept constant in all our experiments (POPC 79.5%; PEG2K 0.5%; Chol 20%), except for GUVs labeled with lissamine Rhodamine PE (Rhod PE), where the composition was (POPC 98%; PEG2K 1%; RhPE 1%). Chloroform solutions containing 0.25 mg of the lipids mix were uniformly spread onto slides coated with indium tin oxide (ITO). The lipid-coated slides were then subjected to a vacuum for over 30 minutes to ensure complete evaporation of the chloroform. A capacitor was assembled by placing a 0.3-mm rubber septum between two lipid-coated slides, and the interspace was filled with approximately 200 μL of a solution containing approximately 350 mM sucrose (∼350 mOsm). Giant unilamellar vesicles (GUVs) with diameters ranging from 10 to 100 μm were electroformed by applying an AC voltage of 1.5 V at 10 Hz across the capacitor for 1 hour following the published protocol (Angelova, M. I. & Dimitrov, D. S. Liposome electroformation. Faraday Discuss. Chem. Soc. 81, 303–311 1986).

#### Recruitment of ssDNA to the GUV surface

Each designed ssDNA synthesized and functionalized by Integrated DNA Technologies (IDT) with a Rhodamine fluorophore at the 5’ end and a cholesterol group at the 3’ end for membrane anchoring. To recruit ssDNA to the GUV surface, a 500 nM ssDNA solution was prepared in PBS. This solution was mixed with the GUV solution at a 1:20 ratio and gently homogenized using a 20 µL micropipette. The mixture was placed on a lab rotisserie and incubated for 10 minutes to prevent precipitation and aggregation of GUVs. The solution was homogenized again using a 20 µL micropipette. This solution was then transferred to a SurFlex imaging chamber (see section Protocol for anisotropy measurement).

### Protocol for anisotropy measurement

#### Preliminary alignment of the excitation polarizer using Rhodamine PE GUVs

Sample preparations for anisotropy measurements are described in the sections above. Prior to anisotropy measurement, the excitation polarizer was aligned in a direction kept constant throughout the experiments: GUVs labeled with RhPE lipids were imaged without analyzer polarizers, and the excitation polarizer was rotated until the maximum signal from the GUV contour was at the top of the image (*SI appendix* Fig. S7A). This ensured that the excitation polarizer orientation remained constant between different experiments. The Hamamatsu Gemini module with PBS and two analyzer polarizers was then inserted to perform the anisotropy measurement.

#### Image acquisition

ssDNA on GUV samples (see section: Recruitment of ssDNA to the GUV surface) were loaded into a custom imaging chamber, made by gluing a p1500 pipette tip to a coverslip. The sample consisted of 5 µL of GUV solution, 100 µL of PBS, and 50 µL of MQ water. The coverslip was previously passivated by incubating it for 1 minute with 10% bovine serum albumin and then rinsed 3 times with buffer (PBS). The sample was then imaged at the equatorial plane of each vesicle, adjusting laser power and exposure time to achieve a signal-to-noise ratio of ∼20. For each sample, a minimum of 30 GUVs were imaged.

### Image analysis protocol for anisotropy calculation

Images taken with the SurFlex present two images of the same object, one showing polarization parallel to the excitation polarization (top image, I_ǁ_), the other perpendicular polarization (bottom image, I_⊥_) see *SI appendix* Fig. S7B. These images were analyzed using a Matlab program described below, which is accessible at the following link (https://github.com/fletchlab-git/Surflex.git).

#### Registration

After opening, images are cropped to separate the two sub-images corresponding to I_ǁ_ and I_⊥_. The program then searches for GUVs present in the I_ǁ_ sub-image using a circle detector (Matlab: “imfindcircles”) and crops the GUV closest to the center of the image. The same procedure is applied to the I_⊥_ sub-image. Next, images corresponding to the I_ǁ_ and I_⊥_ components of the same GUV, are then registered and aligned with each other (Matlab: “imregister” in translation mode).

#### Anisotropy calculation

Anisotropy was then calculated pixel by pixel from the registered I_ǁ_ and I_⊥_ images using equation (2). Anisotropy values were saved as an image.

#### Profile scanning

The image showing the GUV anisotropy was then analyzed by extracting the anisotropy value along the GUV membrane contour. The profiles thus obtained (linescans) were saved as Excel spreadsheets and subsequently fitted using a Mathematica program to extract persistence length values. See section Fitting procedure to extract *L_p_*/*L* for a description of the Mathematica program.

### Calculation of interaction energies with NUPACK 4.0

Using NUPACK 4.0 software, we computed the ensemble of transient structures for each ssDNA oligomer and their associated folding energies. These calculations were performed at 25°C and 100 mM NaCl concentration, matching the conditions used in our anisotropy measurements. For each oligomer, we calculated an interaction energy (-ΔG) obtained by summing the energies of each structure, weighted by their respective probabilities. Specifically, we only considered structures with probabilities greater than 1%.

### Measuring the anisotropy of glycosylated molecules on red blood cells

Single-donor human whole blood (Innovative Research) was used and red blood cells were extracted by centrifugation at 300 G for five min. Red blood cells were washed one time with PBS, and were incubated with labeling lectins (RCA-I fluorescein labeled; Vector Laboratories) at 10 mg/L for 20 min at room temperature. In the case of red blood cells treated with trypsin, the cells were incubated with a Trypsin EDTA 1X solution for either 5 min, 10 min, or 15 min. To stop the cleaving reaction of trypsin, 1:1 FBS was added to quench the enzyme. Then the red blood cells were washed with PBS three times before being labeled with lectins using the same protocol described above. MQ water was subsequently added to reach a dilution of 1:5 PBS to swell cells and ensure that they were spherical. Then cells were imaged in BSA-treated custom-made SurFlex microscopy imaging chamber (see section Image acquisition) and imaged with the protocol described in section “Protocol for anisotropy measurement”.

### Theory Overview

The theoretical framework developed in this work links the optical theory of fluorescence anisotropy with the underlying polymer dynamics. The fluorescence anisotropy r is a powerful technique to infer the physical properties of molecules, such as their orientation and stiffness. The key step in the theory is to express the orientation of the fluorophore absorption and emission dipoles in terms of the angles between them and the polarization of the exciting light (*SI appendix* Eqs. 8-10 in supporting text and Fig. S1). This optical component is then connected to the conformational fluctuations of the tethered polymer, which is modeled using a worm-like chain (WLC) description. The probability distribution function P(θ) for the angle θ between the polymer tip tangent vector and the membrane normal is derived from a 2D WLC model (Eq. 14). This provides a crucial link between the observed anisotropy and the polymer’s persistence length *L_p_*, which characterizes its stiffness. In the stiff limit (*L_p_*/*L* >> 1), an analytical expression for the anisotropy r is obtained (Eq. 15 *SI appendix,* supporting text), which exhibits a *cos*(2*γ*) dependence on the angle *γ* between the membrane normal and the polarization axis of the incident light. Conversely, in the floppy limit (*L_p_*/*L* → 0), the anisotropy takes a simpler form (Eq. 16 *SI appendix,* supporting text) that depends only on the angle β between the absorption and emission dipoles. The derived theoretical expressions provide greater insight into the relationship between the polymer’s physical properties, the fluorophore’s optical characteristics, and the observed fluorescence anisotropy. This integrated theoretical framework sets the stage for the validation and application of the model through molecular dynamics simulations.

### MD simulations overview

To validate the theoretical predictions extensive molecular dynamics (MD) simulations were conducted using the HOOMD-blue software. A coarse-grained bead-spring model was used to represent tethered biomolecules. In the case of single-stranded DNA (ssDNA) 2 beads corresponded to 3 nucleotides. The total potential energy consisted of a Weeks-Chandler-Andersen potential for excluded volume interactions, a Finite Extensible Nonlinear Elastic potential for bonding interactions, and a Kratky-Porod potential for bending rigidity (Eq. 17 *SI appendix,* supporting text). After equilibration, the orientation of the end-tangent vector was characterized by the angle α and compared to the theoretical predictions from the 2D WLC model. Simulations were performed for various persistence lengths *L_p_*, with 32 independent replications. The resulting probability distributions, standard deviations, and maximum percentage errors were analyzed to validate the theoretical framework (*SI appendix* Fig. S2, supporting text). Furthermore, the fluorescence anisotropy r was computed from the MD simulations for different fluorophore lifetimes *τ* and compared to the theoretical predictions (*SI appendix* Fig. S3, supporting text). This allowed for the assessment of the theory’s accuracy in capturing the effect of finite measurement timescales on the experimental observable.

### Fitting procedure to extract *L_p_*/*L*

The fluorescence anisotropy r was expressed in terms of the polymer persistence length *L_p_*, the contour length L, the angle *γ* between the membrane normal and the polarization axis of the incident light, as derived in the theoretical framework (Eq. 15 *SI appendix,* supporting text). To fit the experimental data, we numerically integrated the expression over all the relevant angular degrees of freedom in Mathematica (code accessible at the following link: https://github.com/fletchlab-git/Surflex.git), as described in the *SI appendix,* supporting text. Using the experimental anisotropy value r0=0.374 (corresponding to β=11.9°) from literature (38) for the angle between absorption and emission dipoles, we determined α, the angle of attachment of the fluorophore with respect to the polymer tip tangent vector. This analysis used data from Lissamine-Rhodamine inserted into giant unilamellar vesicles (GUVs) with short linkers, where we obtained Lp/L ≈ 20. After the calibration step, we fitted the experimental data for the single-stranded DNA (ssDNA) conditions individually for each GUV. This was done by using the NonlinearModelFit function in Mathematica to determine the best-fit value of the persistence length *L_p_* for each GUV. The fitting considered the full angular dependence, allowing us to obtain a robust characterization of the ssDNA polymer stiffness for each measurement. The NonlinearModelFit function minimizes the sum of the squared differences between the experimental anisotropy values and theoretical predictions, iteratively updating the model parameter *L_p_* to find the optimal fit. The uncertainty in the fit parameter was found to be negligible compared to the spread in the persistence length values obtained from the individual GUV measurements. Finally, the mean persistence length was calculated by averaging the *L_p_* values obtained from the individual GUV fits, and the standard deviation was computed to reflect the spread in the data. This approach allowed us to leverage the detailed theoretical framework, validated through the molecular dynamics simulations, to quantitatively analyze the experimental fluorescence anisotropy data and extract the physical properties of the ssDNA polymer, i.e the measured *L_p_*/*L* values.

### List of reagents and equipment

#### Reagents

PBS 1X, Corning REF 21-040-CV; BSA 10% (w/w) in MQ water: BSA (Santa Cruz Biotechnology Inc. P/N # SC-2323A); MQ water (Millipore, Milli-Q IQ 7000); POPC, 1-palmitoyl-2-oleoyl-glycero-3-phosphocholine (PN # 850457, Avanti polar lipids); RhodPE: 1,2-dioleoyl-sn-glycero-3-phosphoethanolamine-N-(lissamine rhodamine B sulfonyl) PN # 810150, Avanti polar lipids); Peg 2K PE (PN # 880120, Avanti polar lipids); Cholesterol (PN # 700100, Avanti polar lipids); Lectins RCA1 labeled with fluorescein (Vector laboratories: ref FL-1081); Trypsin: Trypsin EDTA 1X (Corning, ref 25053-CL); FBS (heat-inactivated Fetal Bovine Serum, Gibco ref A38401-01 50 mL); Single donor whole blood (Innovative Research: CAT# IWB1K2E10ML).

#### Hardware

Centrifuge: 5424 Microcentrifuge (cat # 05-400-005 Eppendorf); Thermo scientific lab rotisserie (model M90615 American laboratory trading); glass coverslip (REF 48393-230 24×40mm N 1.5 Avantor); UV glue: Norland optical adhesive 60 (Norland products inc. P/N 5001); Dual view module: W-VIEW GEMINI Image Splitting Optics A12801-01 (Hamamatsu); linear polarizer : LPVISE100-A - Ø1 (Thorlabs inc.); Polarizing beam splitter: WGP00058 FBF04C, 40×25mm, TA=0, VC0 (Moxtek Inc.); rotation stage manual : LRM 1 - Ø1 (Thorlabs inc.); camera: Prime 95B sCMOS camera (Teledyne Photometrics); microscope : Nikon ECLIPSE Ti2 inverted microscope; objective : SR plan Apo IR 60X/ 1.27 WI (Nikon instruments Inc. mat # MRY10060); spinning disk: CSU-X1 Spinning Disk Field Scanning Confocal System (Yokogawa).

Sequence of ssDNAs synthesized by Integrated DNA technology (IDT).

- 20nu: /5RhoR-XN/ATG CGG CGT TGC TAC CGA CG/3CholTEG/
- 30nu: /5RhoR-XN/ATC TGT GTG CGG ACG GTG CGG CGA GTT ACT/3CholTEG/
- 50nu: /5RhoR-XN/TGC GCT ACC TTG CAG GAA TCG AGG CCG TCC GTT AAT TCC CCT TGC GTA CG/3CholTEG/
- polyA: /5RhoR-XN/AAA AAA AAA AAA AAA AAA AAA /3CholTEG/
- polyC:/5RhoR-XN/CCC CCC CCC CCC CCC CCC CCC/3CholTEG/
- CACA:/5RhoR-XN/ACACACACACACACACACACA/3CholTEG
- CCAA: /5RhoR-XN/AAA AAA AAA AAC CCC CCC CCC /3CholTEG/
- AACC: //5RhoR-XN/CC CCC CCC CCA AAA AAA AAA A/3CholTEG
- seq#1: /5RhoR-XN/ GCC CAG GTT ACA GTT CTG CGT/3CholTEG/.
- seq#2: /5RhoR-XN/ATG CTG CGT TGC TAC CGA CGT /3CholTEG/.
- seq#3: /5RhoR-XN/GAT CGT GAG CTT ACT CTC GCG /3CholTEG/.
- CAG repeats: /5RhoR-XN/CAG CAG CAG CAG CAG CAG CAG/3CholTEG/

## Acknowledgments

We would like to thank Dr. Sungmin Son and Dr. Shalin Mehta for early discussions and Fletcher laboratory members for feedback, especially Andrés Dextre for help with NuPack 4.0 and Jared Huzar for advice on DNA oligomer design. D.A.F. is supported by NIH R01 GM134137, the NSF Center for Cellular Construction (DBI-1548297), and the Miller Institute for Basic Research. D.A.F. is a Chan Zuckerberg Biohub Investigator. EMBO Postdoctoral fellowship funded A.C. for this work. S.A. is a Chan Zuckerberg Biohub Collaborative Postdoctoral Fellow. D.A.F. is a Chan Zuckerberg Biohub Investigator.

## Author Contributions

A.C., S.A. and D.A.F. conceived of the study and designed research; A.C. conducted the experiments; L.B. and A.C assayed cells; A.C. and S.A. analyzed the data; S.A. performed molecular dynamics simulations and developed the theory with feedback from A.C; D.A.F. supervised the study; A.C., S.A., and D.A.F. wrote the manuscript; all authors provided feedback on the manuscript.

