## Supporting information for "SurFlex Microscopy: Measuring Flexibility of Surface-Tethered Biomolecules"

#### This PDF file includes:

Supporting text
Figs. S1 to S7
SI References

### Supporting Information Text

### 1. Theory modeling

Measurements of fluorescence anisotropy are a powerful technique to infer the physical properties of polymers, including their orientation and stiffness. When a sample is excited with polarized light, the resulting emission is also polarized; the degree of this emission polarization is described by the anisotropy $r$ (1). In this study, we utilize this technique to directly determine the stiffness, characterized by the persistence length, of tethered polymers, such as those found on cell surfaces.

The experimental setup is illustrated in the main text. Briefly, polarized laser light strikes the fluorophore attached to the end of a polymer tethered to a membrane. The time delay between the absorption and emission of the fluorophore, known as the fluorophore lifetime ($\tau$), typically around 1-10 ns. The quantum energy-level absorption and emission events occur on much faster timescales (order of picoseconds) and are therefore considered instantaneous for our analysis.

During this process, the orientation of the fluorophore changes by an angle $\theta$, which depends on the rotational diffusion time scale of the polymer. For an end-tethered polymer, this time scale is influenced by the chain's stiffness (or persistence length) and the viscosity of the surrounding medium. The emitted polarized light generally has a different orientation from the incident light and is split into two orthogonal directions. One detector measures the intensity of the polarization perpendicular to the incident light, $I_\perp$, while the other detector collects light with the same orientation as the source, $I_\parallel$. This data is collected over longer timescales (approximately microseconds) and thus represents a time-averaged measurement of the polymer chain fluctuations.

**A. Fluorescence Anisotropy Fundamentals.** The anisotropy $r$ of a light source is defined as the ratio of the polarized component to the total intensity ($I_T$),

$$r = \frac{I_z - I_y}{I_x + I_y + I_z}. \quad [1]$$

When the excitation is polarized along the $z$-axis, the emission from the fluorophore is symmetric along that axis. Hence, $I_x = I_y$, $I_y = I_\perp$ and $I_z = I_\parallel$. Thus,

$$r = \frac{I_\parallel - I_\perp}{I_\parallel + 2I_\perp}. \quad [2]$$

We note that the normalization with total intensity, i.e., $I_\parallel + 2I_\perp$ instead of $I_\parallel + I_\perp$ is preferred, since this results in higher sensitivity even when the output signal intensities are low. For a given orientation of the fluorophore, with polar angle $\theta$ and azimuthal angle $\phi$, the parallel and perpendicular intensitiies are given by,

$$I_\parallel = \cos^2\theta, \quad [3a]$$

$$I_\perp = \sin^2\theta \sin^2\phi. \quad [3b]$$

For excitation polarized along the $z$-axis, the excited fluorophore population over time is symmetrically distributed around that axis. Hence, we can eliminate the $\phi$ dependence by averaging and this results in,

$$I_\parallel = \cos^2\theta, \quad [4a]$$

$$I_\perp = \frac{1}{2}\sin^2\theta. \quad [4b]$$

Now we assume that we observe the fluorophore that is oriented relative to the $z$-axis with an excitation probability $f(\theta)$. The measured average fluorescence intensities are,

$$I_\parallel = \int_0^{\pi/2} f(\theta)\cos^2\theta d\theta = k\langle\cos^2\theta\rangle, \quad [5a]$$

$$I_\perp = \frac{1}{2}\int_0^{\pi/2} f(\theta)\sin^2\theta d\theta = \frac{k}{2}\langle\sin^2\theta\rangle, \quad [5b]$$

where we have defined,

$$\langle\cos^2\theta\rangle = \frac{\int_0^{\pi/2} f(\theta)\cos^2\theta d\theta}{\int_0^{\pi/2} f(\theta)d\theta}. \quad [6]$$

Here, $f(\theta)d\theta$ is the probability that the fluorophore is oriented between $\theta$ and $\theta + d\theta$, and $k = \int f(\theta)d\theta$ is a constant that depends on the physical properties of the tethered polymer. Inserting into Eq. (2) results in,

$$r = \frac{3\langle\cos^2\theta\rangle - 1}{2}. \quad [7]$$

Hence the anisotropy is determined by the average value of $\cos^2\theta$, where $\theta$ is the angle of the emission dipole relative to the z-axis. Our task now is to determine the probability $f(\theta)$ for the specific experimental setup that we describe next.

**B. Orientational Degrees of Freedom for Membrane-Tethered Fluorophores.** To calculate the average emitted intensity of fluorophores end-attached to membrane-tethered proteins (specifically giant unilamellar vesicles, GUVs, in our study), it is crucial to express the orientation of the absorption and emission dipoles in terms of the angles between them and the polarization of the exciting light. Since our fluorophore is attached to a protein tethered to the lipid membrane, additional angles must be introduced (see Fig. S1).

We define the polarization direction of the illuminating laser along the x-axis, with the light traveling along the z-axis. The angle between the membrane normal (N) and the polarization axis is denoted as $\gamma$. The orientation of the tip of the membrane-tethered polymer (P) is defined by the azimuthal angle $\theta$, referred to as the Tip Vector Angle (TVA) in the main text, and a twist angle $\tau$. The optical absorption dipole ($f_a$) of the fluorophore is shifted from the polymer tip vector (P) by an angle $\alpha$ and a twist angle $\phi$. In general, the emission dipole ($f_e$) may not be collinear with the absorption dipole, so we introduce an additional rotation by an angle $\beta$ and a twist angle $\delta$.

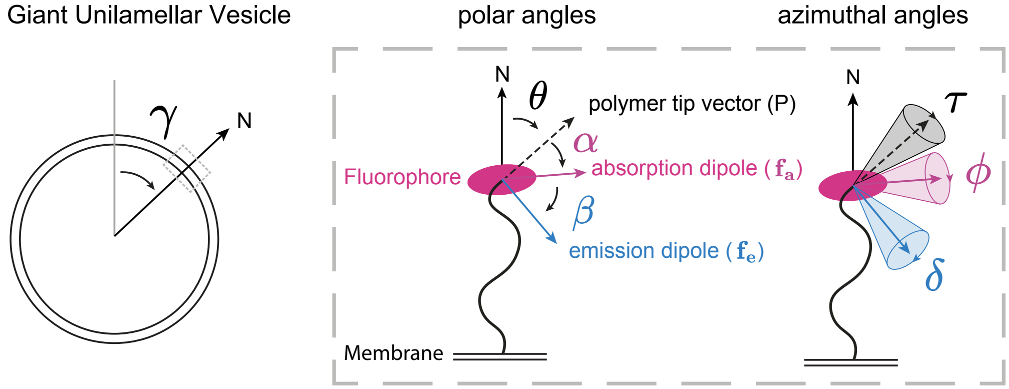


**Fig. S1.** Orientational degrees of freedom of a tethered polymer with a fluorophore at its distal end. The fluorophore absorption (emission) dipole $\mathbf{f_a}$ ($\mathbf{f_e}$) is shifted from its geometric axis $F$, which itself may be non-rigidly tagged on to the tethered polymer, forming a non-zero angle with the polymer tip tangent vector $P$. The polymer is modeled as a worm-like chain and the probability distribution function of its fluctuating end tangent vector relative to membrane normal $N$ is explicitly known. Finally, the membrane normal itself is interrogated along the equatorial plane of the GUV, making an angle $\gamma$ with the incident polarized light dipole (x-axis).

We assume that the membrane normal always lies in the x-y plane. This can be arranged in the experiment by recording confocal microscopy images at the equator of the roughly spherical GUVs. The general absorption dipole vector $\mathbf{f_a}$ can be determined by taking an initial vector (representing the fluorophore optical absorption dipole relative to its geometric orientation) and performing six rotations as illustrated schematically in Fig. S1: a rotation of angle $\phi$ about the x-axis, a rotation of $\psi$ about the z-axis, a rotation of angle $\sigma$ about the x-axis, a rotation of $\alpha$ about the z-axis, a rotation of angle $\tau$ about the x-axis, and finally a rotation of angle $\gamma$ about the z-axis:

$$\mathbf{F_a} = R_z(\gamma)R_x(\tau)R_z(\theta)R_x(\phi)\begin{pmatrix}\cos\alpha\\ \sin\alpha\\ 0\end{pmatrix} \quad [8]$$

Similarly, the emission dipole vector is obtained by performing two additional rotations:

$$\mathbf{F_e} = R_z(\gamma)R_x(\tau)R_z(\theta)R_x(\phi)R_z(\alpha)R_x(\delta)\begin{pmatrix}\cos\beta\\ \sin\beta\\ 0\end{pmatrix} \quad [9]$$

where $\beta$ is the angle between the optical absorption and emission dipoles. Now the angle between the fluorophore absorption (emission) dipole vector and the laser polarization axis, $\Theta_a$ ($\Theta_e$), is given by

$$\cos\Theta_a = \mathbf{F_a}\cdot\mathbf{x}, \quad \cos\Theta_e = \mathbf{F_e}\cdot\mathbf{x}. \quad [10]$$

The averaged quantity required to compute $r$ is the average of $\cos^2\Theta_a\cos^2\Theta_e$ over the angles $\tau$, $\alpha$, $\sigma$, $\phi$, $\theta$, $\delta$, with the appropriate geometric factors included, given by

$$\mathrm{I}_{\parallel} = \langle\cos^2\Theta\rangle = \frac{\int P(\theta)\cos^2\Theta_a\cos^2\Theta_e\sin\theta\, d\delta\, d\theta\, d\phi\, d\sigma\, d\tau\, d\alpha}{\int P(\theta)\cos^2\Theta_a\sin\theta\, d\delta\, d\theta\, d\phi\, d\sigma\, d\psi\, d\tau\, d\alpha} \quad [11]$$

where $\sin\theta$ is the geometric factor to account for spherical coordinates. $P(\theta)$ is the reduced probability density of the polymer tip tangent vector $P$ to have an angular displacement $\alpha$ from the membrane normal $N$. The integration limits of the twist angle span the entire 0 to $2\pi$ interval while the polar angle goes from 0 to $\pi/2$. We will now derive the probability distribution $P(\theta)$ for semi-flexible worm-like chains in the next subsection.

**C. Reduced probability distribution function for worm-like chains.** We consider the specific form for the probability $P(\alpha)$, representing the angle $\alpha$ that the tip tangent vector of a tethered semi-flexible polymer chain makes with the membrane normal $N$. We specialize to a worm-like chain model characterized by a persistence length, $L_p = \frac{EI}{k_B T}$, where $E$ is the Young's modulus and $I$ is the second moment of inertia. We assume that the rotational diffusion time-scale of the tip is much longer than the fluorophore lifetime in the following analysis.

The probability distribution for fluctuations from the mean orientation of the tangent vector at the free end of a tethered worm-like chain (WLC) is given by (2):

$$p(\theta) = \sqrt{\frac{l_p}{2\pi}} \exp\left(-\frac{l_p \theta^2}{2}\right), \tag{12}$$

where $l_p = \frac{L_p}{L}$ is the dimensionless persistence length, and $L$ is the contour length of the semi-flexible polymer, typically determined by its molecular structure. The boundary condition at the tethering point is assumed to be clamped (fixed in both position and angle), which is a reasonable approximation for molecules with transmembrane domains at cell membranes or other rigid interfaces. It is crucial to note that Eq. (12) has been modified from the original reference to account for the difference in persistence length definition between the 2D and 3D WLC models. Specifically, the persistence length in 3D is twice that in 2D for the same polymer stiffness, necessitating a factor of 2 adjustment in the equation. Although Eq. (12) is derived for a 2D worm-like chain (WLC), we argue that it accurately describes the 3D WLC in the stiff regime (persistence length $l_p > 1$. This equivalence between 2D and 3D descriptions arises from the physics of stiff polymers, where thermal fluctuations produce only small deviations from a straight configuration. In the stiff limit, the polymer's behavior is governed by the bending energy, which scales as $(d\theta/ds)^2$, where $\theta$ is the local tangent angle and $s$ is the arc length. The dominant fluctuations are long-wavelength bending modes, which minimize the energetic cost. These fundamental modes are effectively planar—even in 3D space—because out-of-plane deformations require higher spatial derivatives and are thus energetically unfavorable. The polymer's conformational statistics therefore become quasi-two-dimensional. We will verify this theoretical argument through extensive molecular dynamics simulations, comparing predictions from Eq. (12) with full 3D MD trajectories across multiple $l_p$ values spanning the stiff regime.

Next, we consider the distribution of the angles swept out by the fluctuations of the tip tangent vector. Due to the dipolar nature of the emission/detection, there is an inherent periodicity of $\pi$. Therefore, we construct a "reduced" probability distribution that sums the contributions from every $\pi$ rotation:

$$P(\theta) = \sum_{n=-\infty}^{\infty} p(\theta + n\pi). \tag{13}$$

Inserting Eq. (12) into the above expression, the sum converges, and we obtain:

$$P(\theta) = \frac{1}{2\pi} \vartheta_3 \left(\theta, \exp\left(-\frac{1}{2l_p}\right)\right), \tag{14}$$

where $\vartheta_3$ is the Elliptic Theta function of the third kind and is evaluated numerically.

We can explicitly evaluate the anisotropy $r$ by integrating over all angular degrees of freedom in Eq. (11). The problem allows for analytical solutions in two limiting cases: the stiff limit and the floppy limit. In the stiff limit where $l_p > L$, the polymer's orientation remains nearly fixed ($\alpha \approx 0$), allowing us to derive:

$$r = \frac{1}{16}(1 + 3\cos 2\beta)(1 + 3\cos 2\gamma) \tag{15}$$

where $\gamma$ is the angle between the membrane normal and the polarization axis of the incident light. Although our 2D WLC expression was derived assuming $l_p > 1$, we can still analyze the floppy limit ($l_p \ll 1$) because the reduced probability distribution $P(\theta)$ approaches a constant value as $l_p \to 0$. This physical behavior correctly captures how a completely flexible polymer can explore all tip angles with equal probability, matching the expected orientational decorrelation in 3D. In this regime, we obtain:

$$r = \frac{2}{5}\left(\frac{3\cos^2\beta - 1}{2}\right) \tag{16}$$

where $\beta$ is the angle between absorption and emission dipoles. This result matches Lakowicz (1) and is independent of $\gamma$, reflecting the complete orientational averaging of the fluorophores.

The persistence length $l_p$ strongly affects the measured anisotropy $r$. For floppy chains ($l_p \ll 1$), thermal fluctuations cause the polymer to rapidly explore all possible conformations, leading to orientational averaging of the fluorophore dipoles and consequently low anisotropy values. Conversely, stiff chains ($l_p > 1$) maintain their orientational correlation along the contour, restricting the fluorophore motion and resulting in higher anisotropy values. This direct relationship between polymer stiffness and fluorescence anisotropy enables the experimental determination of persistence length through polarization measurements.

 Aymeric Chorlay[1], Siddhansh Agarwal[1,2], Lena Blackmon[1], Daniel Fletcher[1,2,3]

### 2. Molecular Dynamics Simulations

To validate our theoretical predictions and explore the applicability of the 2D WLC model to 3D systems in the stiff limit, we conducted extensive molecular dynamics simulations using HOOMD-blue (3). Our approach employed a coarse-grained bead-spring model to represent single-stranded DNA (ssDNA), striking a balance between computational efficiency and physical accuracy.

**A. Model Description.** We model ssDNA using a discretized worm-like chain framework, with coarse-grained dimensions to capture the flexibility and dynamics of the molecule. In this model, the ssDNA molecule is represented as a chain of spheres with a radius of 0.5 nm. Given that the average spacing between nucleotides in ssDNA is approximately 0.63 nm, each pair of spheres in our coarse-grained model represents the spatial extent of about three nucleotides (4). While in experiments we vary the contour length (number of bases) to effectively change $L_p/L$, in our simulations we fix the chain length at $N = 20$ beads and directly vary the bending rigidity $\kappa$ to access different $L_p/L$ ratios. This approach maintains consistent numerical accuracy across all stiffness regimes while capturing the same essential physics of the experimental system, as the polymer behavior is governed by the dimensionless ratio $L_p/L$ rather than the absolute values of $L_p$ or $L$.

The total potential energy of our system was given by:

$$U_{\text{total}} = U_{\text{WCA}} + U_{\text{FENE}} + U_{\text{bend}} \qquad [17]$$

where $U_{\text{WCA}}$ represents the Weeks-Chandler-Andersen potential for excluded volume interactions, $U_{\text{FENE}}$ is the Finite Extensible Nonlinear Elastic potential for bonding interactions, and $U_{\text{bend}}$ is the Kratky-Porod potential for bending rigidity.

***A.1. Excluded Volume Interactions.*** We modeled excluded volume interactions using the WCA potential, a shifted and truncated version of the Lennard-Jones potential that is purely repulsive:

$$U_{\text{WCA}}(r) = \begin{cases} 4\epsilon\left[\left(\frac{\sigma}{r}\right)^{12} - \left(\frac{\sigma}{r}\right)^{6}\right] + \epsilon, \text{if } r < 2^{1/6}\sigma \\ 0, \text{if } r \geq 2^{1/6}\sigma \end{cases} \qquad [18]$$

We set $\epsilon = 1.0$ (in units of $k_BT$) and $\sigma = 1.0$. This choice ensures a strong short-range repulsion between beads, effectively implementing the excluded volume interaction without any long-range attraction.

***A.2. Bonding Interactions.*** To model the covalent bonds between adjacent beads, we employed the FENE potential:

$$U_{\text{FENE}}(r) = -0.5kr_0^2 \ln\left[1 - \left(\frac{r}{r_0}\right)^2\right] \qquad [19]$$

Here, we chose $k = 20.0$ and $r_0 = 1.5$, values that have been shown to prevent bond crossing while allowing for realistic bond fluctuations (5).

***A.3. Bending Rigidity.*** To incorporate the semiflexible nature of ssDNA, we implemented bending rigidity through the Kratky-Porod potential:

$$U_{\text{bend}}(\theta) = \kappa(1 - \cos\theta) \qquad [20]$$

where $\theta$ is the angle between adjacent bonds, and $\kappa = L_p k_B T$, with $L_p$ being the persistence length in simulation units. This formulation allowed us to easily adjust the stiffness of our polymer by varying $L_p$.

**B. Simulation Protocol.** We implemented a cantilever boundary condition by fixing the positions of the first two beads of the polymer chain. This clamped-end constraint reflects how ssDNA interacts with the lipid membrane of GUVs, where the insertion of the attachment chemistry into the bilayer can restrict rotational motion at the anchoring point. This provides a reasonable starting approximation for modeling the polymer dynamics in our system. The system was equilibrated using Langevin dynamics at constant temperature ($k_BT = 1.0$) for $10^6$ timesteps. We employed a timestep of 0.005 in reduced units, striking a balance between simulation accuracy and computational efficiency.

To relate our simulation timescales to physical time, we consider the translational diffusion of a single bead in water near a solid surface. For a single bead with radius $a \approx 0.5$ nm, the bulk translational diffusion coefficient $D_0$ can be estimated using the Stokes-Einstein relation: $D_0 = k_BT/(6\pi\eta a)$, where $k_B$ is Boltzmann's constant, $T = 293$ K (room temperature), and $\eta \approx 1$ mPa·s (the viscosity of water at room temperature). Substituting these values, we obtain $D_0 \approx 10^{-10}\text{m}^2/\text{s}$. The presence of the membrane significantly modifies the local hydrodynamics. First, the no-slip boundary condition at the membrane surface creates a position-dependent friction tensor that scales as $1/h$, where $h$ is the distance from the surface. Second, the tethering constraint restricts the accessible conformational space. These effects combine to reduce the effective diffusion coefficient. Experimental and computational studies of polymer segmental dynamics near interfaces show a reduction in mobility by approximately two to three orders of magnitude (6–11). Taking this into account, we estimate $D_{\text{effective}} = D_0/100 \approx 10^{-12}\text{m}^2/\text{s}$ for a tethered polymer segment near the membrane surface.

Using this effective diffusion coefficient, we can map the simulation timescale to physical time. In our simulations, one time unit corresponds to a distance traversed by diffusion over a characteristic length scale $a \sim 0.5$ nm, thus the real time per simulation time unit is $t_{\text{real}} = a^2/D_{\text{effective}} \approx 250\,\text{ns}$. This calculation implies that each simulation timestep corresponds to approximately 1.25 ns of physical time. The application of the slowdown factor here accounts for the substantial reduction in diffusion expected near an interface, aligning well with the experimental observations in systems with moderate grafting densities.

**C. Data Collection and Analysis.** After equilibration, we computed the tangent vector at the free end of the polymer as the unit vector pointing from the penultimate to the last bead. We characterized its orientation by the angle $\alpha$ from the xz plane, storing this data every 1000 timesteps.

We performed simulations for fixed chain length ($N = 20$) and different persistence lengths, with each condition replicated 32 times using different random seeds to ensure robust statistics. The resulting distributions of $\alpha$ were then compared to the predictions of Eq. (12), allowing us to validate the applicability of the 2D WLC model in the stiff limit of 3D WLC. This comparison, detailed in the following section, provides crucial insights into the dimensional crossover of polymer behavior in the stiff regime.

### 3. Results and Discussion

Figure S2 presents a comprehensive comparison between our 3D molecular dynamics simulations and the theoretical predictions derived from the 2D WLC model.

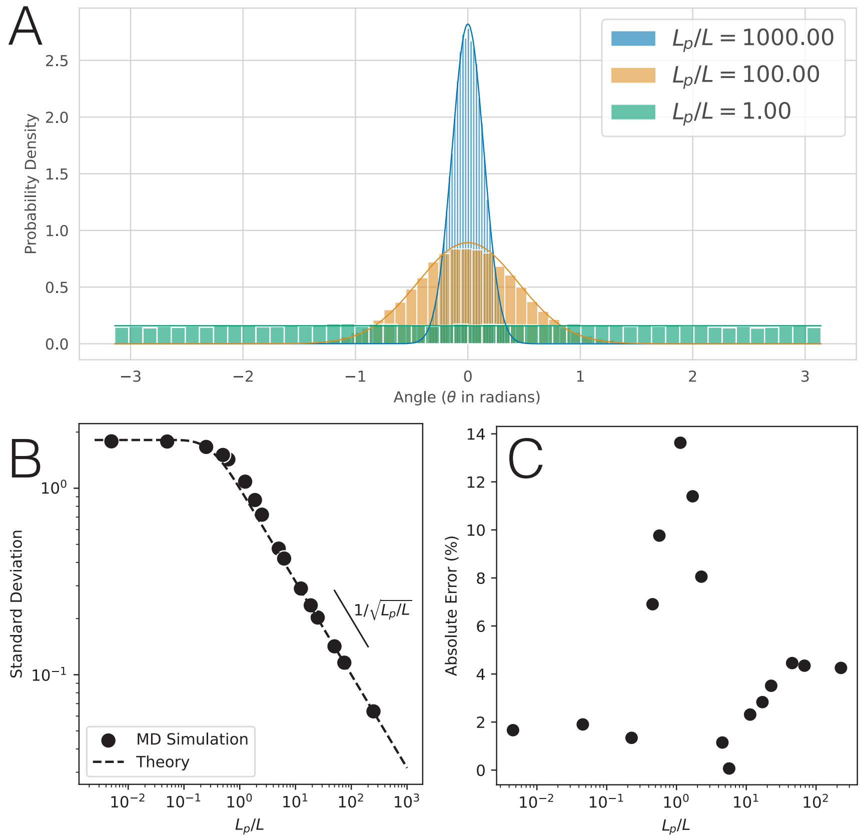


**Fig. S2.** Comparison of 3D molecular dynamics simulations with 2D WLC theory. (A) Histogram of probability density (normalized to 1) from MD simulations (bars) overlaid with the theoretical prediction from Eq. (14) (solid curve) for various $l_p$ ratios. (B) Standard deviation of the end-tangent angle $\alpha$ as a function of $l_p$, comparing MD results (points) with theoretical predictions (line). (C) Maximum percentage error between MD simulations and theoretical predictions across different $l_p$ values.

Panel A demonstrates the remarkable agreement between the probability density distribution of the end-tangent angle $\alpha$ obtained from our MD simulations and the theoretical prediction given by Eq. (14). The histogram bars represent the MD data, while the solid curves show the theoretical distributions. This agreement holds across various $l_p$ ratios, indicating the robustness of our 2D WLC approximation in describing 3D polymer behavior in the stiff regime.

In panel B, we plot the standard deviation of $\alpha$ as a function of $l_p$. The MD simulation results (points) closely follow the theoretical prediction (line) derived from 14. This quantitative agreement further validates the applicability of our 2D WLC model to 3D systems in the stiff limit.

Panel C quantifies the discrepancy between simulation and theory by showing the maximum percentage error across different $l_p$ values. Notably, the error remains below $\sim 15\%$ for all studied stiffness regimes. Our 2D WLC approximation is effective in capturing the essential physics of 3D semiflexible polymers, particularly when $l_p > 1$, the regime of interest in this study.

 Aymeric Chorlay[1], Siddhansh Agarwal[1,2], Lena Blackmon[1], Daniel Fletcher[1,2,3]

The close agreement between 3D MD simulations and 2D WLC theory supports our hypothesis that as $l_p$ increases, the polymer's configurational space becomes increasingly confined, which suppresses out-of-plane coiling. This allows the 2D model to accurately describe the 3D system's behavior. The minimal discrepancy observed, particularly at intermediate $l_p$ values, suggests that our approach captures the dominant physical mechanisms governing the polymer's end-tangent orientation. However, the slight increase in error at very high $l_p$ ratios may indicate the onset of rod-like behavior, where the assumptions underlying the WLC model begin to break down.

### 4. Effect of fluorophore lifetime

To further validate our model and explore its implications for experimental observables, we computed the fluorescence anisotropy from our MD simulations for various fluorophore lifetimes and compared these results with theoretical predictions.

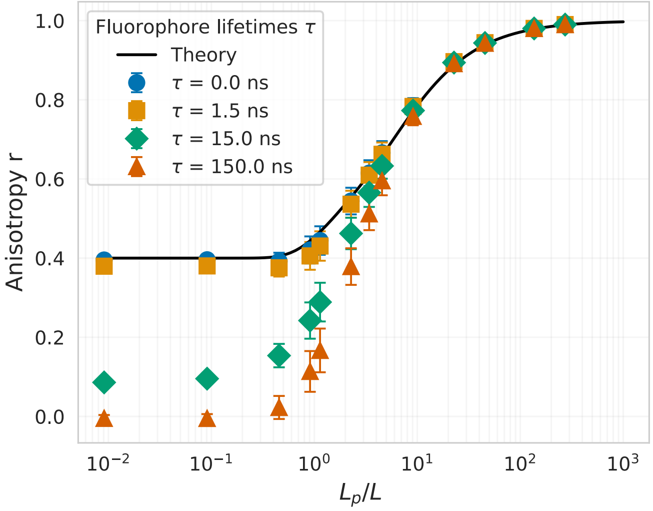


**Fig. S3.** Fluorescence anisotropy as a function of $l_p$ for different fluorophore lifetimes. Solid line represents the theoretical prediction. Symbols show MD simulation results for fluorophore lifetimes $\tau = 0.0$ ns (instantaneous), 1.5 ns, 15.0 ns, and 150.0 ns. Error bars represent standard deviation from 32 independent simulations.

Figure S3 presents the fluorescence anisotropy $r$ as a function of the normalized persistence length $l_p$ for various fluorophore lifetimes $\tau$. The solid line represents the theoretical prediction derived from our 2D WLC model, while symbols denote results from MD simulations. To obtain these results, we performed 32 independent simulations for each $l_p$ value, collecting time-correlated data on the orientation of the end-tangent vector. Specifically, for each MD trajectory, we computed the time-correlated fluorescence anisotropy by evaluating:

$$r(\tau) = \frac{3\langle \cos^2 \Theta_a(t) \cos^2 \Theta_e(t+\tau)\rangle - 1}{2} \quad [21]$$

where $\Theta_a(t)$ and $\Theta_e(t+\tau)$ are the angles between the excitation polarization axis and the absorption and emission dipole orientations at times $t$ and $t+\tau$, respectively. The averaging $\langle\cdot\rangle$ was performed over all possible time origins $t$ and across all 32 independent trajectories. Error bars represent the standard deviation across trajectories. The anisotropy was then computed using time lags corresponding to different fluorophore lifetimes ($\tau = 0.0$ ns, 1.5 ns, 15.0 ns, and 150.0 ns).

The data reveal excellent agreement between theory and simulation for fluorophore lifetimes on the order of a few nanoseconds—such as those typical of Rhodamine dyes used in this study (12)—across the entire range of $l_p$ values studied. This agreement is particularly strong in the stiff regime ($L_p/L > 1$), corroborating our earlier findings on the applicability of the 2D WLC model to 3D systems in this limit. For longer fluorophore lifetimes (15.0 ns and 150.0 ns), we observe increasing deviation from the theoretical curve, especially at lower $l_p$ values. This discrepancy can be attributed to the fact that our theory assumes instantaneous measurement of the end-tangent orientation, while longer fluorophore lifetimes allow for reorientation of the polymer during the fluorescence emission process. Notably, all curves converge in the high $l_p$ limit, approaching the theoretical maximum anisotropy of 1. For very stiff polymers, the end-tangent orientation remains essentially static over the timescales of fluorescence emission, regardless of the fluorophore lifetime.

However, in the flexible regime ($L_p/L < 1$), selecting an optimal fluorophore lifetime becomes more nuanced. For polymers with high flexibility, a fluorophore lifetime that is too short may not fully capture segmental dynamics, while one that is too long risks averaging out critical orientation fluctuations. There is thus a "sweet spot" for the fluorophore lifetime, where it is long enough to register meaningful fluctuations in the polymer orientation but short enough to avoid significant reorientation during emission. Identifying this optimal lifetime as a function of polymer flexibility presents an interesting direction for future studies.

### 5. Experimental Data Fitting Procedure

Our fitting procedure for the experimental fluorescence anisotropy data incorporates an additional angular parameter $\theta_{\text{sys}}$ to account for systematic optical alignment of the experimental setup. This parameter represents a rotation between the excitation polarizer and the parallel analyzer, which manifests as a modification of the reference axis for measuring polarized intensities. In practice, this requires modifying the reference vector in Eq. (10) from $(1, 0, 0)$ to $(\cos\theta_{\text{sys}}, \sin\theta_{\text{sys}}, 0)$ when computing the angles between the fluorophore absorption/emission dipoles and the polarization axis.

The numerical integration over all angular degrees of freedom was performed in Mathematica, incorporating this additional rotation matrix. Across all experimental measurements, we consistently obtained $\theta_{\text{sys}} \approx 50°$, independent of the specific sample conditions. This systematic angle affects the absolute values of the measured anisotropy but, importantly, does not alter the amplitude of the anisotropy variations with membrane normal orientation, which remains determined by the polymer's $l_p$ ratio.

The fundamental anisotropy $r_0$ represents the maximum theoretical anisotropy for a fluorophore in the absence of any rotational motion, determined solely by the angular difference between its absorption and emission dipole moments $\beta$. This angle $\beta$ relates to $r_0$ through:

$$r_0 = 0.4\left(\frac{3\cos^2\beta - 1}{2}\right) \qquad [22]$$

where 0.4 is the limiting anisotropy for parallel absorption and emission dipoles ($\beta = 0$). For Rhodamine-based dyes, $r_0 = 0.374$ from literature (13, 14), and this yields $\beta = 11.9°$. Using this value, we determined the angle of attachment $\alpha$ of the fluorophore with respect to the polymer tip tangent vector through fitting of data from Lissamine-Rhodamine in GUVs with short linkers ($L_p/L \approx 20$). The best fit value of $\alpha \approx 90°$ is consistent with the molecular structure of Lissamine-Rhodamine.

For fitting the experimental data, we used the complete numerical solution that integrates over all orientational degrees of freedom, rather than the analytical approximations valid in limiting cases. For each experimental condition, we fitted the anisotropy data from individual GUVs using Mathematica's NonlinearModelFit function with $l_p$ as the sole fitting parameter. The reported persistence lengths represent the mean of these individual measurements, with error bars indicating the standard deviation across multiple GUVs (n $\geq$ 10 for each condition).

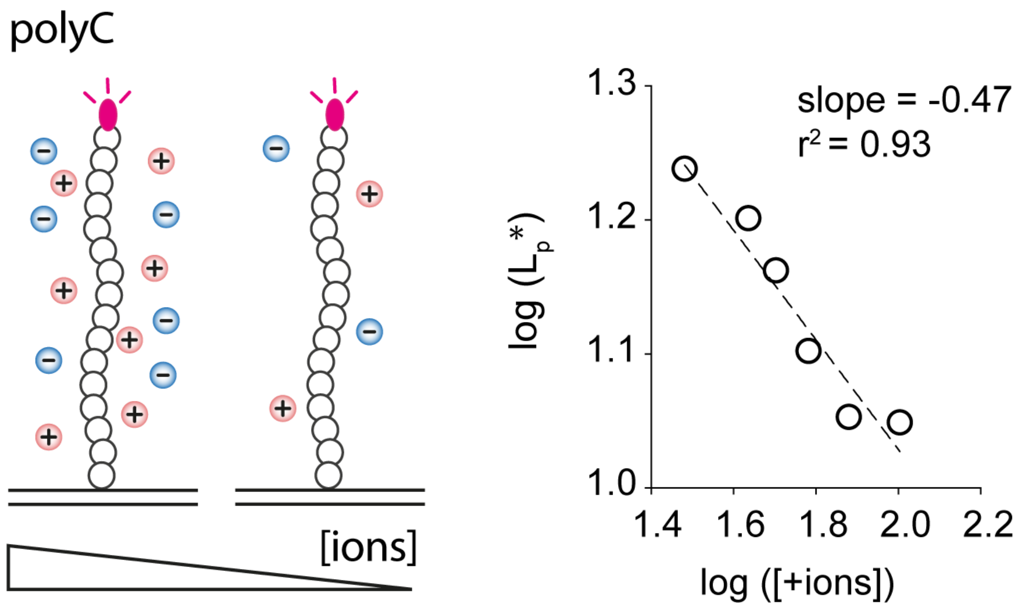


**Fig. S4. Effect of ionic strength on PolyC apparent persistence length and log-log linear regression. Left**: Measurement of the apparent persistence length ($L_p^*$) of polyC (21 cytosine) ssDNA oligomers while decreasing the ion concentration in solution by adding MQ water. **Right**: Log-Log representation of the data from figure 3B : Apparent persistence length ($L_p^*$) as function of positive ion concentration. Dash line show linear regression fit. Slope is 0.47 and the r2=0.94 suggesting that the $L_p^*$ scale as [+ions]$^{1/2}$ as postulated in Joanny et al. (15).

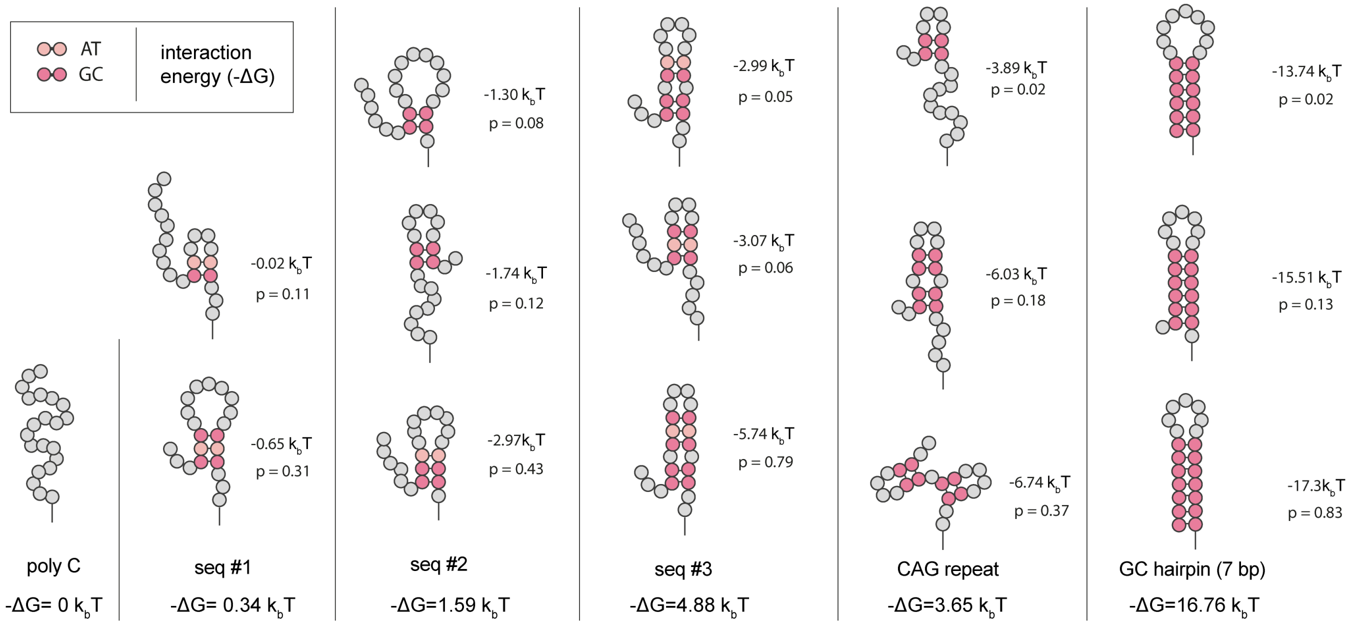


**Fig. S5. Structures and associated energies for ssDNA used in Figure 4.** Graphical representation of the most probable structures of the ssDNA oligomers (studied in Figure 4) and associated energies (in $k_B T$ units) obtained with Nupak 4.0 software. p is the probability of each structure. Bottom line values show the associated interaction energy (-$\Delta$G) calculated by weighting the individual energies of the three most prevalent structures by their respective probabilities. A-T base paires are highlighted in light pink and G-C base paires that have greater binding energy, are highlighted in darker pink.

 

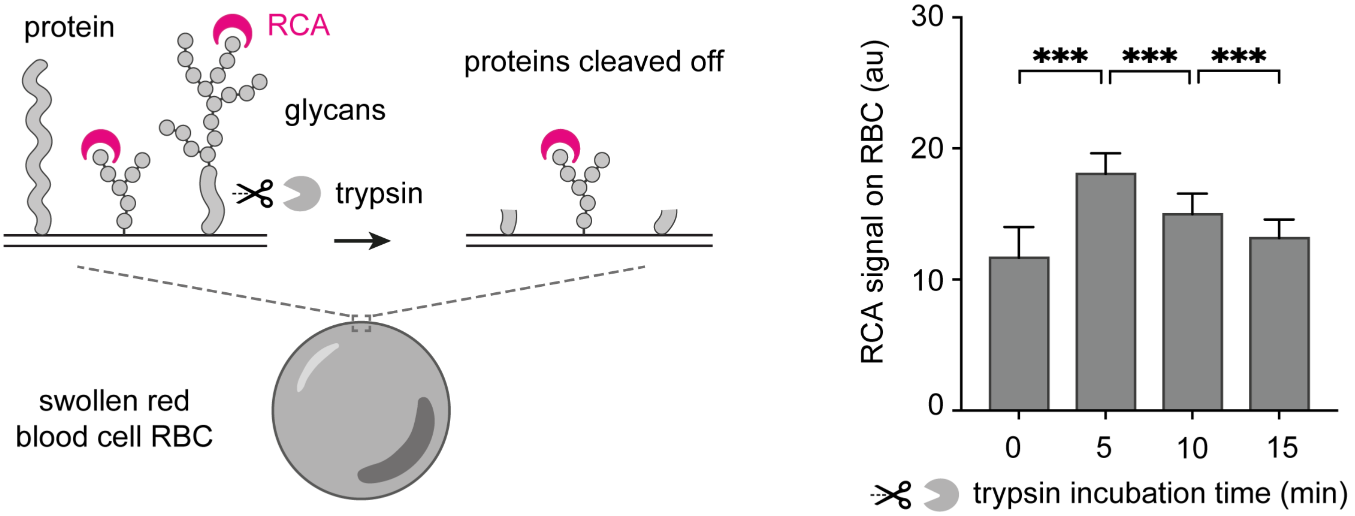


**Fig. S6. Lectin (RCA-I) recruitment level on red blood cells during trypsinization. Left**: Osmotically swollen Red blood cells (RBCs). The terminal galactose residues of glycosylated molecules on the RBC surface are labeled with fluorescein-conjugated RCA lectin. Surface proteins on the RBCs are cleaved by incubation with trypsin. **Right**: Signal of fluorescein-labeled RCA lectin on the surface of RBCs incubated with trypsin for 0 to 15 minutes. Data were obtained by measuring the RCA signal of n=30 independent RBCs for each condition. P-values are < 0.0001 based on t-tests. Error bar indicates standard deviation.

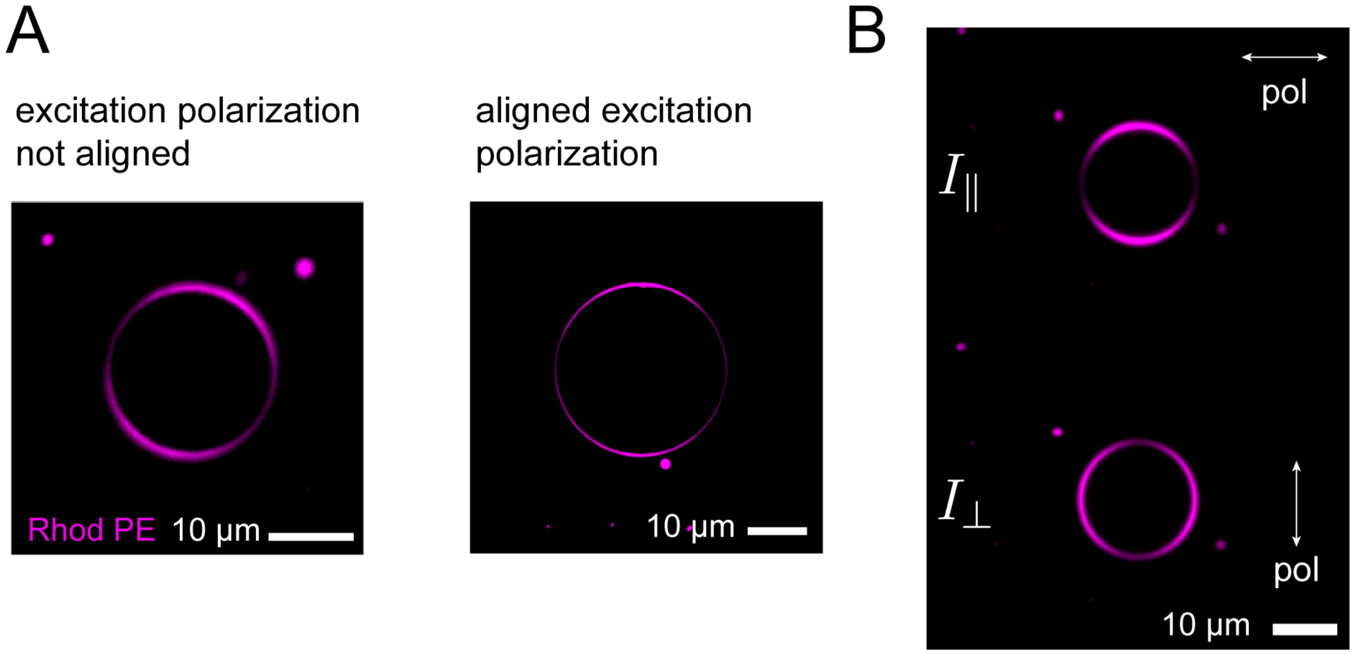


**Fig. S7. Excitation polarization alignment and image analysis illustrations (A)** Illustration of alignment procedure using GUV containing Rhodamine PE lipid (Rhod PE). Left is an image of a GUV when the excitation polarizer is not aligned. Right is an image of a GUV when the excitation polarizer is aligned. **(B)** Raw image of a single GUV containing Rhodamine PE lipid that the Matlab program analyses. Top image: light intensity polarized parallel to the excitation polarization ($I_{||}$); Bottom image: light intensity polarized perpendicular to the excitation direction ($I_{\perp}$). Arrows indicate the orientation of the the two analyzer polarizers.

 Aymeric Chorlay[1], Siddhansh Agarwal[1,2], Lena Blackmon[1], Daniel Fletcher[1,2,3]
